# Distribution patterns of invasive alien plant species in mainland Portugal

**DOI:** 10.1101/2025.06.27.661711

**Authors:** Raquel Fernandes, Andry Castro, Hélia Marchante, Elizabete Marchante, César Capinha

**Affiliations:** Centre of Geographical Studies, Institute of Geography and Spatial Planning, University of Lisbon, Rua Branca Edmée Marques, 1600-276 Lisboa, Portugal; Research Center for Natural Resources, Environment and Society (CERNAS), Polytechnic University of Coimbra, Coimbra Agriculture School, Bencanta, 3045-601 Coimbra, Portugal; Centre for Functional Ecology – Science for People & the Planet, Department of Life Sciences, University of Coimbra, Calçada Martim de Freitas, 3000-456 Coimbra, Portugal; Associate Laboratory Terra, Portugal

**Keywords:** Baseline distribution, drivers of invasion, invasion atlas, occupancy, species richness

## Abstract

Understanding the distribution and diversity of invasive alien species is increasingly vital to meet legal requirements and guide effective management. Portugal currently hosts a large number of invasive alien plant species, with significant environmental and socio-economic impacts. However, the distribution of most of these species in the territory and the factors driving their spread and diversity remain largely unexplored. To address this, we present the first atlas of invasive plants in mainland Portugal. A total of 96 terrestrial and aquatic invasive plants are represented, encompassing all species listed under Portuguese national legislation and the European Union’s list of invasive species of concern. Occurrence data were collected from a broad array of sources, including national and international biodiversity observation databases, citizen science data, literature, and data collections owned by managers and researchers, totaling approximately 85,000 records (mean = 879 ± 1,757 per species). Each species was characterized based on multiple distributional parameters, and a k-means analysis was used to group species with similar distribution patterns. The richness of invasive plants was mapped at the municipality level, and the drivers of their spatial variation were investigated using a comprehensive set of 30 variables representing multiple environmental and human factors. Results show that invasive plants occur in all mainland Portuguese municipalities, but with high variability. Four main patterns of distribution were identified: species that are primarily located along the coast (e.g., *Acacia saligna*), and a gradient of species with narrow (e.g., *Reynoutria japonica*), moderate (e.g., *Ipomoea indica*), and wide distribution ranges (e.g., *Cortaderia selloana*). Species richness was significantly higher in coastal and larger municipalities, particularly those closer to major urban centers and with a higher density of power lines. Our results provide the first comprehensive assessment of the distribution of invasive alien plants in mainland Portugal, establishing a much-needed baseline for future invasion prevention and management efforts.

## Introduction

Human activities are facilitating the global movement and introduction of species beyond their native ranges (Pyšek et al. 2020; Lopez et al. 2022; Capinha et al. 2023). Currently, over 37,000 established alien species have been documented worldwide (IPBES 2023), with approximately 200 new alien species (i.e., ‘exotic’ or ‘non-native’, *sensu* Essl et al. 2019) recorded annually. Among these, plants constitute a significant proportion, with more than 13,000 alien plant species established outside their natural ranges (Seebens et al. 2023).

A substantial portion of alien plants become invasive, causing significant and diverse negative impacts in regions of introduction. These impacts include reductions in native species diversity and abundance (Powell et al. 2011; Hansen et al. 2020; Beaury et al. 2022), disruption of key ecosystem services (Vieites-Blanco and González-Prieto 2020; Ferreira et al. 2021; Ren et al. 2021; Gallardo et al. 2024), and major economic losses affecting multiple activities, including the loss of crops and productive land (Richardson and Wilgen 2004; Maitre et al. 2011; Milanović et al. 2020; Reynolds et al. 2020; Ruiz and Carlton 2003; Sirbu et al. 2022). Additionally, several invasive plants pose environmental risks to human health (Rodriguez et al. 2021). Most of these impactful species have been reported in the American continent, Asia-Pacific region, Central Asia, and Europe (IPBES 2023). In Europe, at least 7,335 alien vascular plant species have been recorded, of which 1,037 (about 14%) are considered invasive in at least one European territory (Kalusová et al. 2024).

Given the frequent severity of the observed and potential impacts, many national and regional governments now prioritize managing biological invasions, including such species in dedicated legislation. Portugal was one of the first European countries to adopt legislation on invasive alien species (hereafter invasive species), introducing in 1999 a list of species subject to specific restrictions, including bans on commercial use and possession (Ministério do Ambiente 1999). This list included over 40 plant species, with several listed as invasive such as *Acacia dealbata, Carpobrotus edulis,* and *Oxalis pes-caprae*, and others considered with known ecological risk (either in the early stages of invasion or having a high potential to become invasive in the future), such as *Ludwigia peploides, Reynoutria japonica,* and *Senecio inaequidens*. This legislation and associated species list were updated by Decree-Law No. 92/2019 (Ministério do Ambiente e Transição Energética 2019), which now includes more than 100 plant species listed as invasive for mainland Portugal. The updated list includes confirmed and potential invasive species known to occur in the territory, as well as others that are presumed absent but expected to become invasive if introduced. Additionally, the European Union Regulation No. 1143/2014 (Official Journal of the European Union 2014) includes the “Union list”, which specifically lists the invasive species of Union concern. This regulation aims to prevent and mitigate negative environmental impacts on European Union countries. Species listed are automatically part of the Portuguese national list, being subject to Decree-Law No. 92/2019. As of the third update of the Union list in 2022, 41 plant species are listed (Official Journal of the European Union 2022).

Over the years, a relevant number of studies have investigated plant invasions in Portugal (Sousa et al. 2018). For example, several have listed the diversity of alien plants (Almeida and Freitas 2000; Almeida and Freitas 2012; Marchante et al. 2014), distribution or potential invasion risk (Rodríguez-Merino 2023). Others have analyzed individual species such as *Baccharis spicata* (Verloove et al. 2017), *Carpobrotus edulis* (Maltez-Mouro et al. 2009; Chefaoui and Chozas 2018), *Cotula coronopifolia* (Costa et al. 2009), or *Pontederia crassipes* (Pádua, Antão-Geraldes, et al. 2022; Pádua, Duarte, et al. 2022; Mouta et al. 2023). Studies have also focused on *Acacia* species (Marchante et al. 2003; Martins et al. 2016; Vicente et al. 2019; Raposo et al. 2021; Große-Stoltenberg et al. 2023), a highly problematic group of taxa, with nine species already established in the territory (Marchante et al., 2023). Specific regions and ecosystems have also received particular attention, including protected areas (Gutierres et al., 2011; Duarte et al., 2023), dune habitats (Duarte et al., 2023), forests (Ferreira et al., 2021), riparian systems (Bernez et al., 2006; Aguiar and Ferreira, 2013; Pabst et al., 2022) and inland waters (Oficialdegui et al. 2023).

Despite the numerous valuable contributions and growing existence of citizen science distribution records in different platforms (e.g., project “Invasoras.pt” at iNaturalist), the overall distribution of invasive plants across mainland Portugal remains largely unknown. This gap persists despite national inventories of invasive species being essential tools for monitoring invasions, guiding policy, and formulating action plans (Latombe et al. 2017; Pabst et al. 2022; Sirbu et al. 2022). The lack of comprehensive distribution data is likely due, in part, to the information on individual taxa being dispersed across numerous sources, making collation and mapping a resource-intensive and challenging task. Additionally, for many species, information about the presence of species at local scales is incomplete or missing.

Here, we present a comprehensive atlas of the distribution of invasive plant richness in mainland Portugal. This atlas was created by compiling and harmonizing data from a wide range of sources, including national and international biodiversity observation databases, scientific literature, and data collections from managers and researchers. In Portugal, municipalities play a central role in territorial management, environmental planning, and decision-making regarding field interventions. This is also the case for local and national-level resource allocation for invasive species prevention and management, citizen-science initiatives, and public awareness campaigns. Hence, municipalities were selected as reference sampling units. With information collected, we proposed to: *i*) map the recorded distribution of each invasive alien plant species in the territory; *ii)* identify species sharing similar pattern of distribution; *iii*) assess spatial patterns of species richness; and *iv*) assess the environmental and socio-economic drivers of spatial variation in species richness values. Our results are expected to provide substantial support for decision-making in local-to-national scale prevention and management efforts of alien plant invasions in Portugal.

## Materials and methods

### Study area

Mainland Portugal is located in the western Iberian Peninsula, at the southwestern edge of Europe (Fig. 1). Its territory spans both the Atlantic and Mediterranean biogeographical regions (Roekaerts 2002). The central and southern (Alentejo and Algarve) regions are characterized by a Mediterranean climate with dry, hot summers, while the northern regions experience a temperate oceanic climate with dry, mild summers (Beck et al. 2023). The southern and coastal areas are generally low-lying, whereas the northern and central regions are more mountainous, with several ranges exceeding 1,000 meters in elevation (Ramos et al. 2020). The territory is divided into 278 municipalities. The population is primarily concentrated in coastal municipalities, particularly in the metropolitan areas of Lisbon and Oporto (Meneses et al. 2017). Forests occupy large areas in all Territorial units for Statistics II (NUTS II), especially in the Centre region, while agricultural areas are most represented in Alentejo and the Lisbon metropolitan area (Alves et al. 2022).

**Figure 1.**
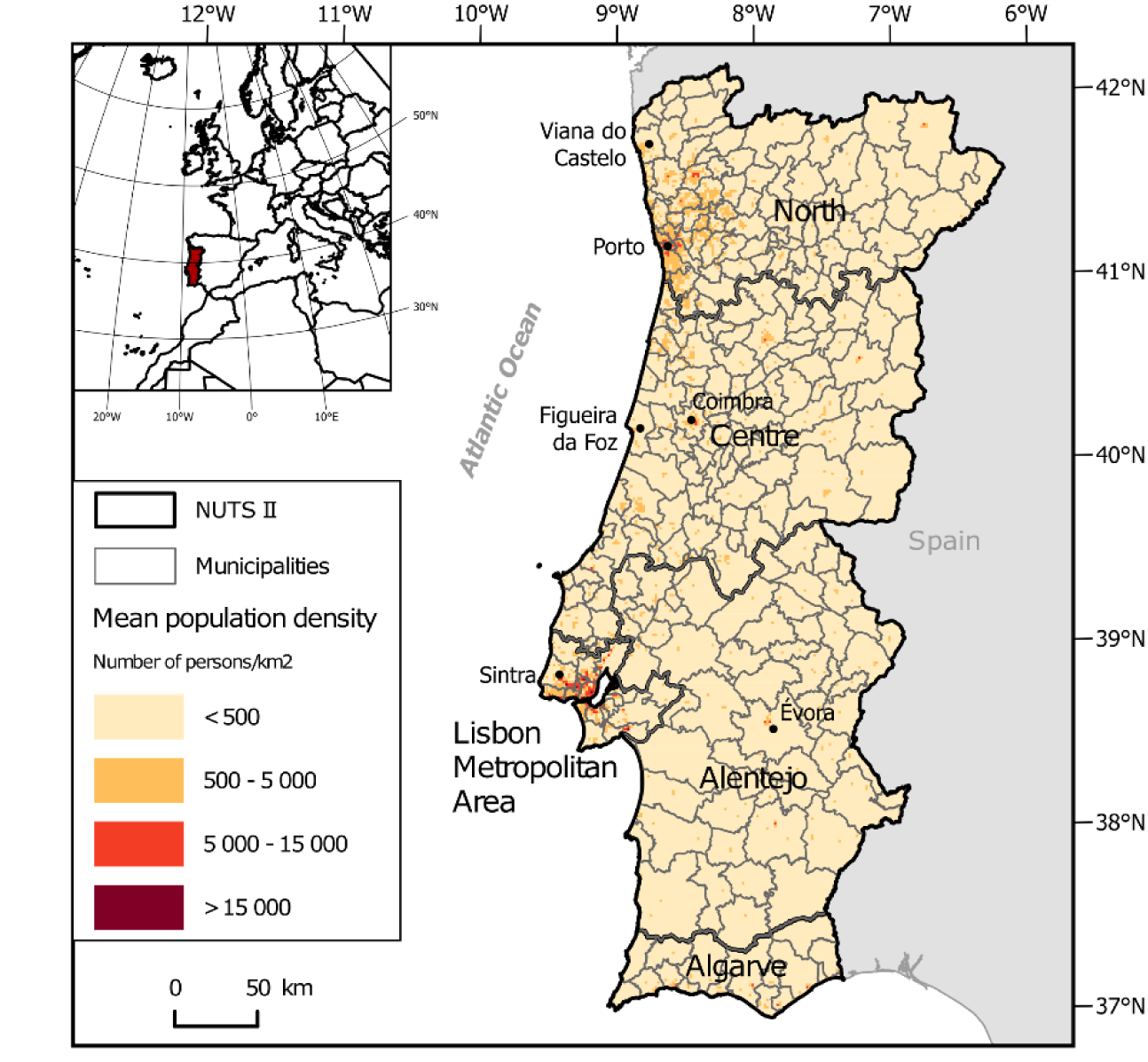
Location of mainland Portugal, showing its administrative divisions, including NUTS II regions (outlined in black), and the 278 municipalities (outlined in grey), used as units of analysis in this study. Population density (persons/km²) is highest in the Lisbon and Oporto Metropolitan Areas, represented by dark red regions.

### List of invasive plants

The mapping effort was focused on the plant species listed in the Portuguese Decree-Law No. 92/2019 (Ministério do Ambiente e Transição Energética 2019) and the European Regulation No. 1143/2014 (Official Journal of the European Union 2014). Since our focus is on mainland Portugal, species mentioned in national legislation that pertain only to the insular regions, such as the Madeira archipelago, were excluded. Portuguese Decree-Law No. 92/2019 lists the genus *Acacia* collectively, unlike other taxa, which are listed at the species level. Therefore, we consulted relevant literature (Marchante et al. 2014, 2023) to identify the *Acacia* species known to occur in the territory. Additionally, we included *Watsonia meriana* in our mapping, as the research community recognizes its invasive behavior in several areas in mainland Portugal (Marchante et al. 2014; Morais et al. 2017). A total of 96 species from 64 genera were included in our inventory, since it was possible to obtain distribution data (Suppl. material 1: Table S1.1).

### Occurrence database

We gathered occurrence records for each invasive plant from multiple sources. The online databases consulted included the Global Biodiversity Information Facility (GBIF) (GBIF.org 2024), Flora-On (Flora-On: Flora de Portugal Interactiva 2024 - an expert-led project aimed at systematizing information for all vascular plant species in Portugal, including their distribution - and the citizen science platforms, iNaturalist (iNaturalist 2024) (of which only research-grade records were considered), and Pl@ntnet (Pl@ntNet 2024). Records were also collected from numerous research publications identified in the “Web of Science” and “Scopus”, combining the search terms “Portugal” and each species name, and from master’s and doctoral theses identified from “RENATES”, a Portuguese platform containing official information on dissertations produced in Portugal (DGEEC 2024). Additionally, unpublished data from the authors, collaborators, and companies that manage invasive plants were also included (see “Acknowledgments” section). Data until May 2024 was considered.

The taxonomy of species was defined and harmonized according to the GBIF backbone taxonomy (GBIF Secretariat 2023). The occurrence records were collected for all available years and included subspecies (if existing). Records identified as having coordinates of stored specimens (e.g., herbarium locations) rather than the actual place of observation in the wild were removed. Additionally, records with a spatial precision of less than 100 meters were excluded.

An exception was made for data from Flora-On, as this source is crucial for the comprehensiveness of the mapping but provides data as centroids of grid squares at a 10×10 km resolution. Despite its contribution to spatial comprehensiveness, the lower geographical precision may introduce some spatial errors in attributing occurrences to municipalities (the spatial units of mapping and analysis used, see below). Therefore, we performed two mapping exercises and associated analyses: one including Flora-On data and another without it. The results from both datasets are highly congruent (see Results section). Thus, we primarily describe the results, including the entire set of occurrence data, while those excluding Flora-On data are provided in the Suppl. material 3.

Given the breadth of consulted sources, we expect to have obtained a representative identification of the species occurring in each municipality. However, as with any similar exercise, our data may still exhibit some gaps and biases, particularly towards regions with higher recording efforts and species that are more conspicuous and easier to identify (more details on the Discussion).

### Invasive plant richness mapping and occupancy

To analyse species distribution patterns, we aggregated occurrence records at the municipality level, following the boundaries defined by the Carta Administrativa Oficial de Portugal (CAOP) (Direção-Geral do Território, 2024). For each municipality, duplicate records of the same species were removed, retaining only one occurrence per species. Species richness was then calculated as the total number of unique species recorded per municipality. To assess species occupancy patterns, we quantified the number of 1×1 km grid cells within each municipality where a species was recorded. The mean percentage of occupied grid cells across all municipalities was then computed to represent overall species occupancy. All analyses were performed using R v4.3.2 (R Core Team, 2022).

### Assessing patterns of species distributions

To identify similarities in species-level patterns of distribution across the territory, we first characterized the distribution of each species using five variables: the number of municipalities where the species occur, the median and interquartile range of distance to the coast (from the centroid of each municipality, in kilometers), and the median and interquartile range of latitude (in decimal degrees).

We then performed a cluster analysis based on these variables. To account for multicollinearity among the variables, we used Principal Component Analysis (PCA), which explained 72.3% of the variance with the first two components (Suppl. material 1: Fig. S1.1). Next, we performed a k-means clustering analysis based on the scores of each species in the bi-dimensional space of the PCA. The PCA was performed using the *’prcomp’* function from the ‘stats’ R package. The k-means procedure employed the *’kmeans’* function from the same package, and the elbow method (Bholowalia and Kumar 2014) was used to identify the optimal number of clusters (Suppl. material 1: Fig. S1.2).

We also examined whether the number of municipalities where a species occurs reflects its overall occupancy across the territory. To do this, we fitted a linear model relating the mean occupancy of species (i.e., the average percentage of occupied 1×1 km grid cells per municipality) to the total number of municipalities where it was recorded.

### Spatial drivers of invasive plants richness

#### Explanatory variables

To identify the factors driving the spatial variation in the richness of invasive plants across the territory we selected a comprehensive set of 30 variables (Table 1) representing distinct environmental and human-related factors and including proxies for recording effort. These variables were mapped at the municipality level and provide a comprehensive representation of the hypotheses on driving factors considered in previous studies (Vicente et al. 2010; Santos et al. 2011; Vicente et al. 2019; Vieites-Blanco and González-Prieto 2020, Pabst et al. 2022).

**Table 1.**
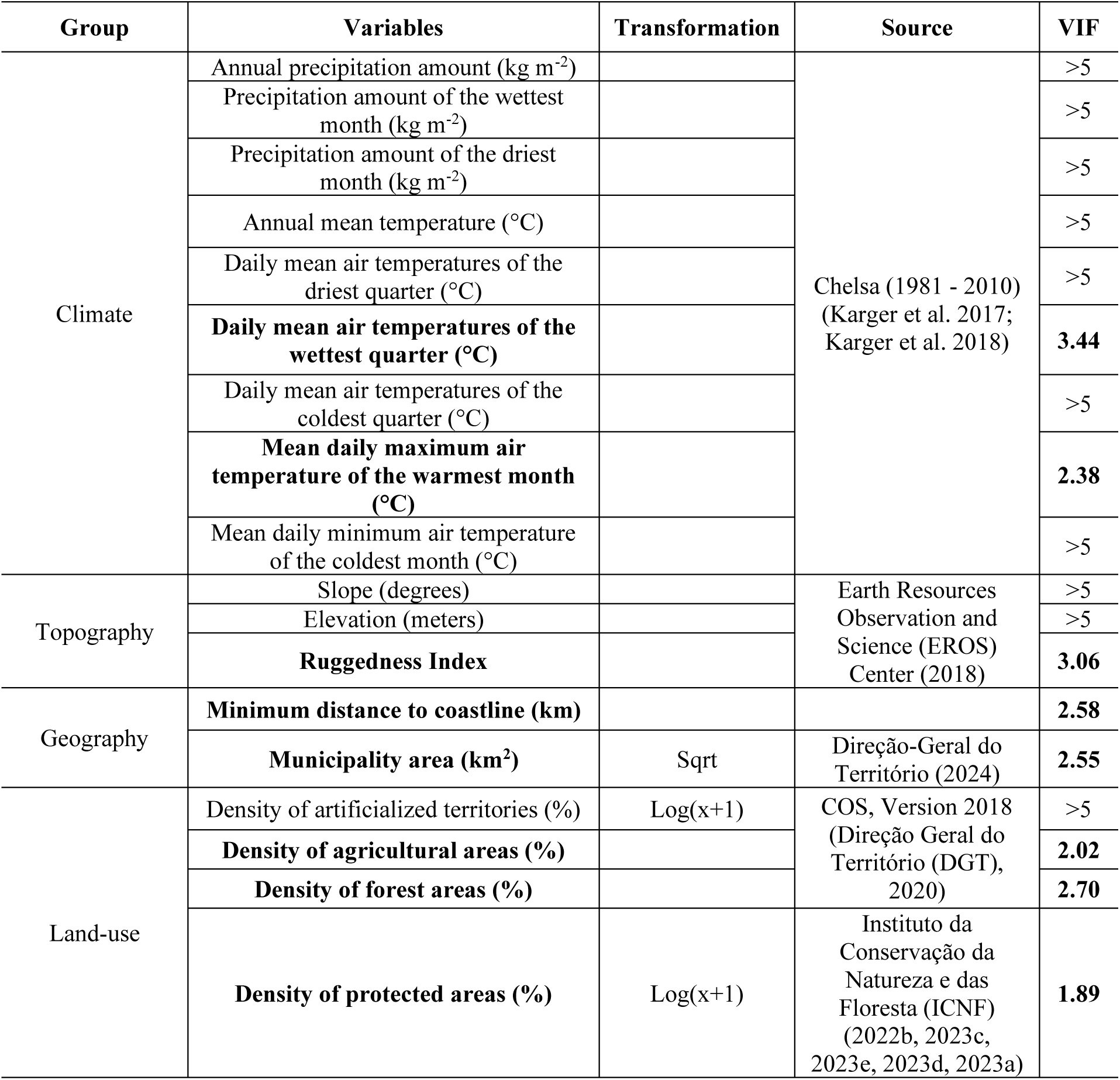
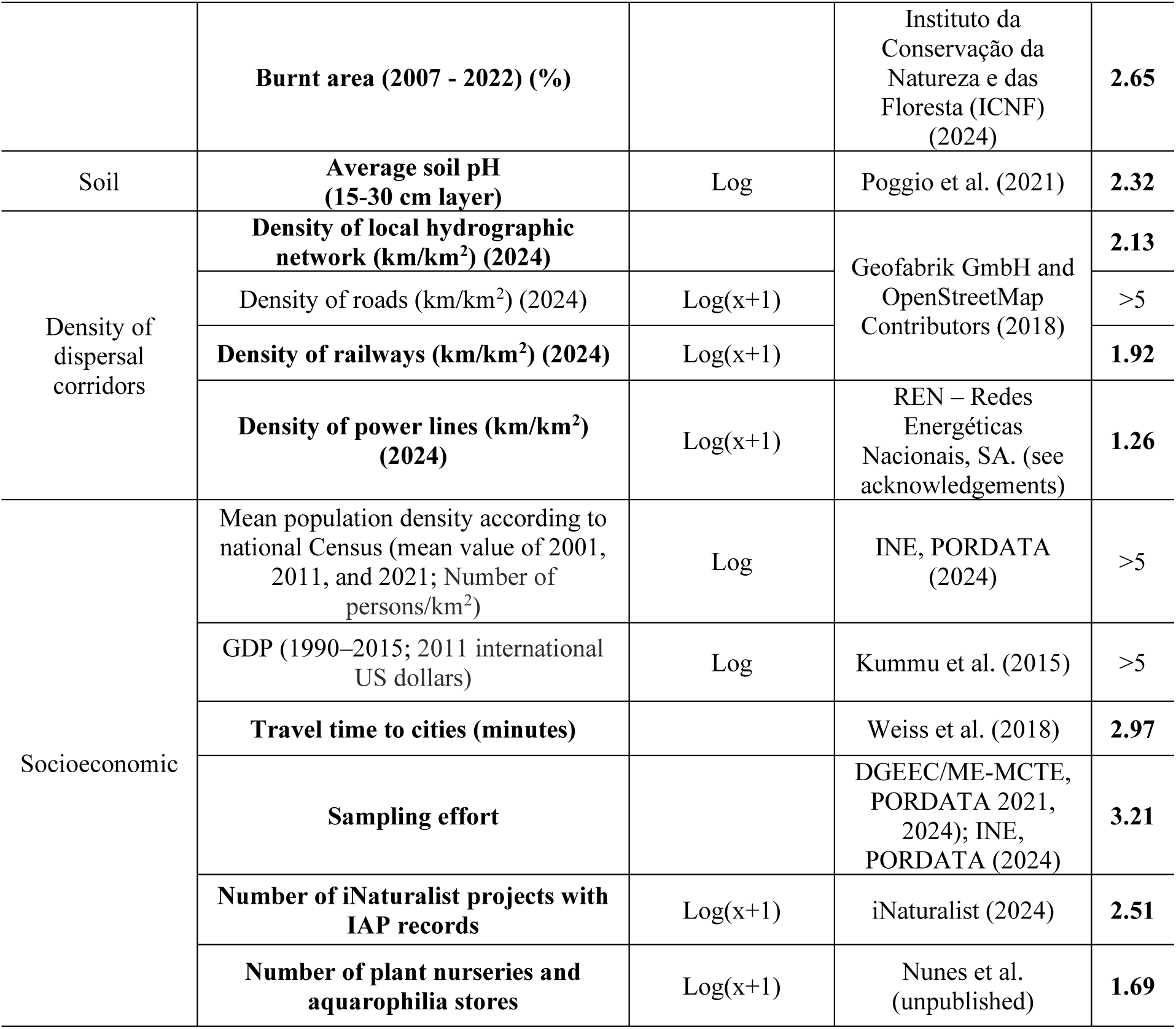
Explanatory variables considered as predictors of spatial variation patterns of invasive plants richness. The final set of variables (in bold) was determined after assessing multicollinearity.

The variables representing climatic conditions, topography, continentality, socio-economy (GDP and travel time to cities, i.e., the time it takes to travel to the nearest urban center via surface transport) (Weiss et al. 2018), and soil groups consisted of mean values of raster cell data within the boundary of each municipality. The land-use variables used vector-type base data and were obtained by dividing the total area occupied by each land-use class by the municipality’s total area. Additionally, variables representing dispersal corridors were calculated by summing the extension of these entities (hydrographic network, road infrastructure, railways, and power lines) within each municipality, divided by its area. The minimum distance to the coastline was determined as the shortest distance from the centroid of each municipality.

The availability of species distribution data can be strongly mediated by varying levels of recording effort (Tiago et al. 2017). Therefore, we considered a set of variables expected to represent spatial patterns in this effort across the territory. These variables were: the average number of higher education students per municipality, the number of higher education institutions per municipality, and the number of iNaturalist projects with invasive plants records per municipality. To avoid redundancy in their spatial variation, we projected these variables into a PCA. The first principal component, explaining around 75% of the variance, was retained, representing mainly the joint variation of the number of higher education students and institutions. The scores of this first composite were extracted and used as a new variable named “academic sampling effort” (Table 1). Because the number of iNaturalist projects contributed little to this component, we used the original values of this variable as a proxy for “citizen-science sampling effort”.

#### Testing for drivers of invasive plants richness

To assess the relationship between the richness of invasive plants across the territory (dependent variable) and the explanatory variables (Table 1), we used a generalized least squares (GLS) regression model (Dormann et al. 2007). Prior to model fitting, square root, and logarithmic transformations were applied to several explanatory variables (Table 1). The response variable was also log-transformed to meet model expectations of a Gaussian distribution of error (Ives 2015).

Potential multicollinearity issues among the explanatory variables were assessed using the variance inflation factor (VIF) (Alin 2010), calculated with the *’vifstep’* function from the ‘usdm’ R package (Naimi 2022). Only the set of explanatory variables with a VIF < 5 were considered for the GLS model (Table 1).

This model was fitted using the *’gls’* function from the ‘nlme’ R package (Pinheiro et al. 2023), assuming an exponential decay in the spatial autocorrelation of richness values (Dormann et al. 2007). Significant relationships between dependent and explanatory variables were determined at ɑ = 0.05, and the goodness-of-fit of the model was measured using the pseudo-R^2^ of Nakagawa et al. (2017).

### Results

#### Invasive plants over Portuguese municipalities

Of the 96 invasive plants with distribution records in mainland Portugal (Suppl. material 1: Table S1.1), 86% are terrestrial, with a smaller portion being aquatic - 14% - including amphibian plants such as *Alternanthera philoxeroides* and *Ludwigia* spp.. They include grasses and herbs (47%), trees and shrubs (29%), and only a few climbing plants (5%) and other growth forms (Suppl. material 1: Fig. S1.3). These species belong to 36 plant families, mainly *Asteraceae* (16%), *Amaranthaceae* (14%), *Fabaceae* (14%), and *Poaceae* (9%) (Suppl. material 1: Fig. S1.4).

Of the ten species with records in more municipalities, the majority are grasses and herbs (50%, e.g., *Datura stramonium, Cortaderia selloana*) and trees and shrubs (40%, e.g.,

*Acacia dealbata, Ailanthus altissima*) (Table 2). Six are already present in more than 200 municipalities (Table 2). The species present in only a few municipalities (up to 10, Suppl. material 1: Table S1.2) are mostly aquatic (40%, e.g., *Nymphaea mexicana* or *Ludwigia grandiflora*). Four species have been recorded in a single municipality (*Alternanthera philoxeroides, Amaranthus hypochondriacus, Amaranthus muricatus,* and *Baccharis halimifolia*, Suppl. material 1: Table S1.2). The distribution of each species per municipality is provided in the Suppl. material 2.

**Table 2.** The ten invasive plants recorded in the greatest number of Portuguese municipalities.

| Species | Number of municipalities with records of the species |
| --- | --- |
| *Acacia dealbata* | 249 |
| *Datura stramonium* | 233 |
| *Arundo donax* | 232 |
| *Phytolacca americana* | 224 |
| *Oxalis pes-caprae* | 213 |
| *Cortaderia selloana* | 208 |
| *Ailanthus altissima* | 198 |
| *Acacia melanoxylon* | 190 |
| *Agave americana* | 187 |
| *Robinia pseudoacacia* | 186 |

The three species with the highest mean percentage of occupancy in mainland Portugal — *Oxalis pes-caprae* (7.53%), *Arundo donax* (6.90%), and *Cortaderia selloana* (6.87%) — are also some of the most widely distributed, occurring in more than 200 municipalities (Table 2; Suppl. material 1: Table S1.3). Their broad geographic range reflects both their high local presence and widespread dispersal across the country. In contrast, *Arctotheca calendula* (5.88%) and *Acacia longifolia* (5.58%) rank fourth and fifth in mean occupancy but have a more restricted national distribution. Their presence is concentrated in coastal regions, suggesting high local presence but limited expansion at a broader scale (Suppl. material 1: Table S1.3).

Similarly, the regression model assessing the relationship between the number of municipalities in which a species was recorded and its mean occupancy revealed a statistically significant positive association (p < 0.05), explaining 42% of the observed variation. This suggests that species occurring in a greater number of municipalities also tend to occupy larger areas within those municipalities. However, the substantial unexplained variation indicates that additional factors influence species occupancy patterns (Fig. 2).

**Figure 2.**
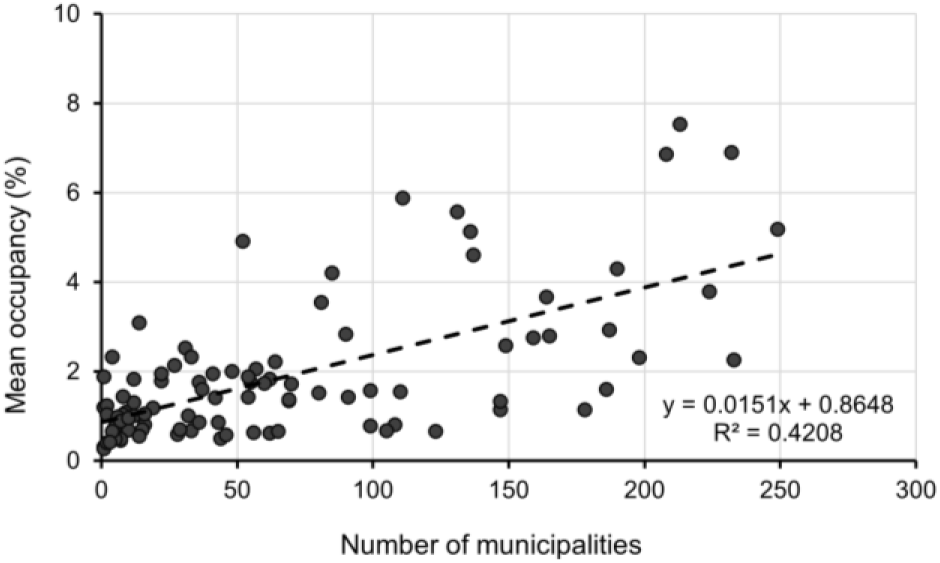
Linear regression model showing the relationship between the total number of municipalities where each species was recorded and its average occupancy (%). Each dot represents a species. The dashed line indicates the linear relationship fitted.

#### Distribution patterns of invasive plants richness

In terms of their distribution patterns, species are grouped overall in four distinct clusters (Fig. 3, Suppl. material 1: Table S1.4 and Suppl. material 3: Fig. S3.2). The first cluster (Fig. 3) includes 26 species predominantly recorded in coastal areas, such as *Acacia saligna* (Fig. 4a) and *Cotula coronopifolia*. The remaining three groups are differentiated mainly by the breadth of their distribution ranges. The second cluster includes 14 species characterized by a narrow distribution area (up to 33 municipalities). These species are also generally found primarily in central and northern municipalities (e.g., *Reynoutria japonica*; Fig. 4b and *Baccharis halimifolia*). The third cluster includes 29 mostly moderately widespread species (e.g., *Acacia longifolia* and *Ipomoea indica*; Fig. 4c). Although there are 10 species with narrow distribution (up to 33 municipalities, Suppl. material 1: Table S1.4), they were grouped in the third cluster due to their distribution over a wider latitude range than the species in the second cluster. The final group consists of 27 species, mostly with very wide ranges (mean number of municipalities = 145 ± 67). Examples of species included in this last group are *Acacia dealbata* and *Cortaderia selloana* (Fig. 4d).

**Figure 3.**
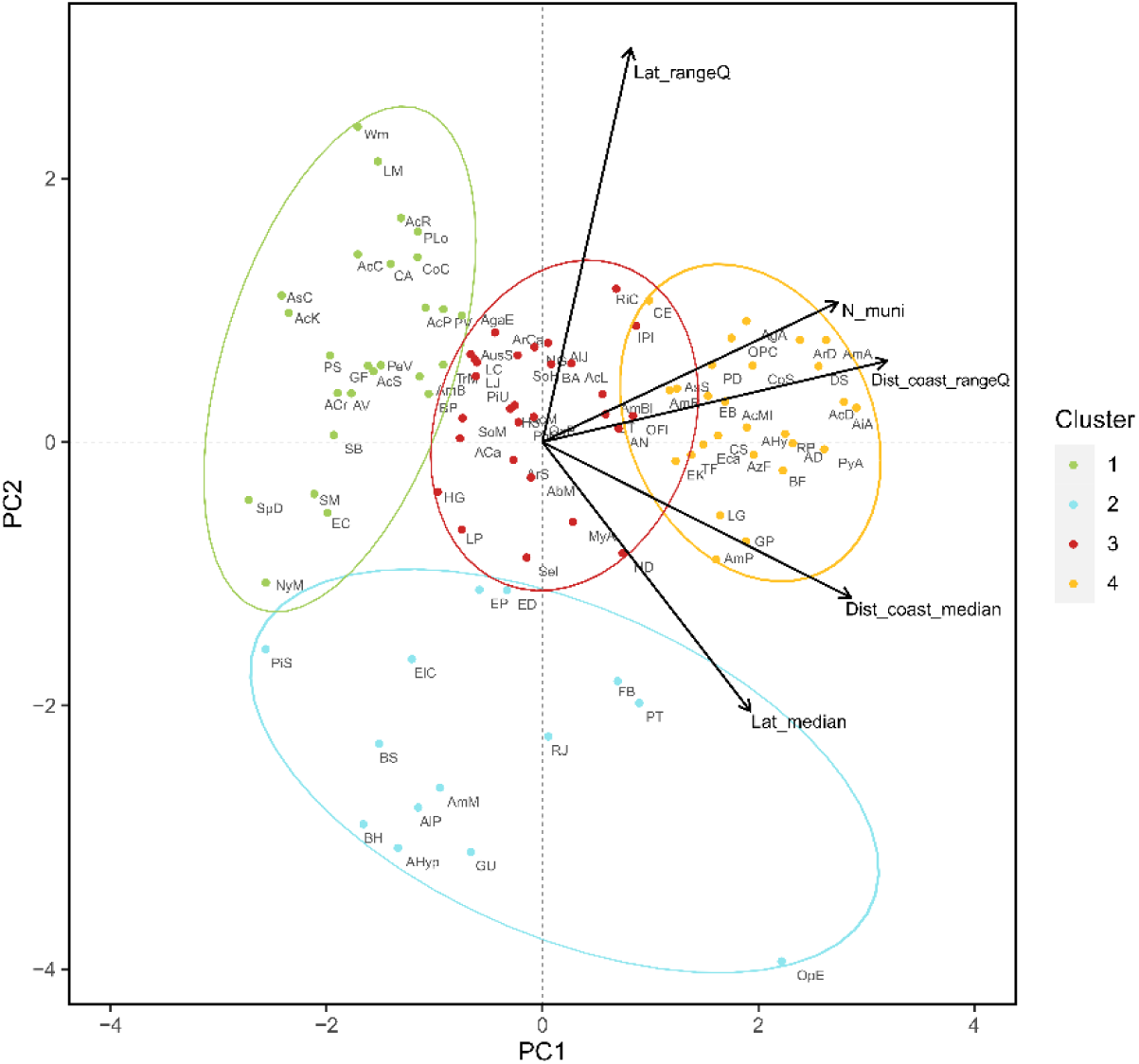
Clusters of invasive plants according to their distribution patterns in mainland Portugal. Cluster 1 (green): species primarily located along the coast; Cluster 2 (blue): mostly species with narrow ranges; Cluster 3 (red): moderately widespread species; Cluster 4 (yellow): mostly widespread species. The two PCA axes represent variation in total number of municipalities of occurrence (N_muni), the median distances to the coast (Dist_coast_median), the interquartile range of distances to the coast (Dist_coast_rangeQ), the median latitudes (Lat_median), and the interquartile range of the latitudes (Lat_rangeQ).

**Figure 4.**
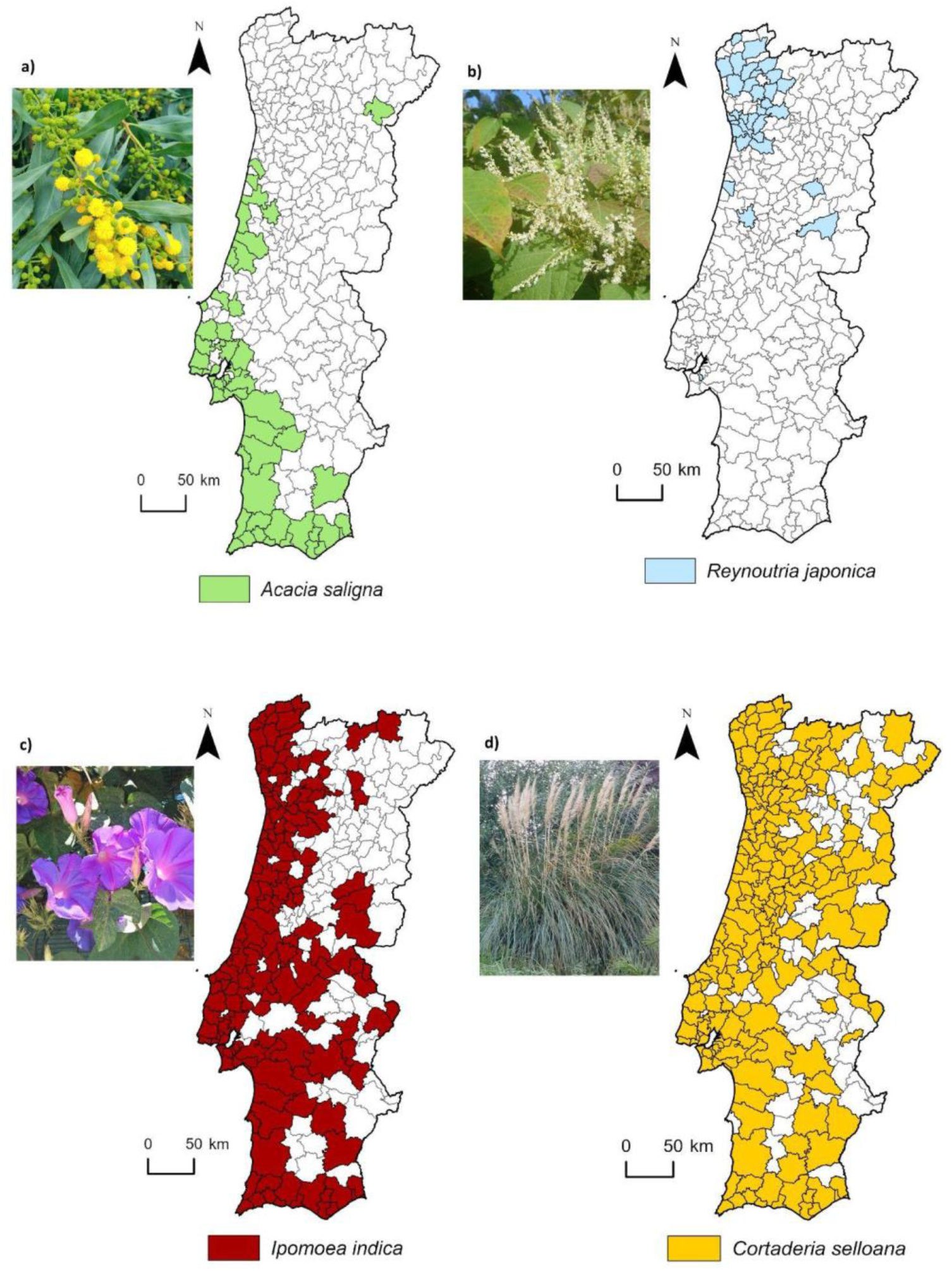
Distribution of example species within each cluster, grouped according to the k-means clustering (Fig. 3), along the Portuguese territory: a) cluster 1 (green) - *Acacia saligna*; b) cluster 2 (blue) - *Reynoutria japonica*; c) cluster 3 (dark red) - *Ipomoea indica*; d) cluster 4 - *Cortaderia selloana*.

#### Patterns and drivers of invasive plants richness

The map of species richness of invasive plants (Fig. 5, Suppl. material 3: Fig. S3.1) shows that at least one invasive plant species has been recorded in each municipality. The number of species is markedly higher in coastal municipalities, particularly those near major urban centers such as the metropolitan areas of Lisbon and Oporto. Additionally, certain cities of regional significance, including Coimbra, Figueira da Foz, and Viana do Castelo, as well as at locations along the Algarve coast in southern Portugal, also exhibit high numbers of invasive plants. The highest numbers were recorded in Lisbon (63 species), Coimbra (62), Sintra (58), and Figueira da Foz (55). Conversely, lower species richness was observed in municipalities primarily located in northeastern Portugal and some central and inland regions of Alentejo.

**Figure 5.**
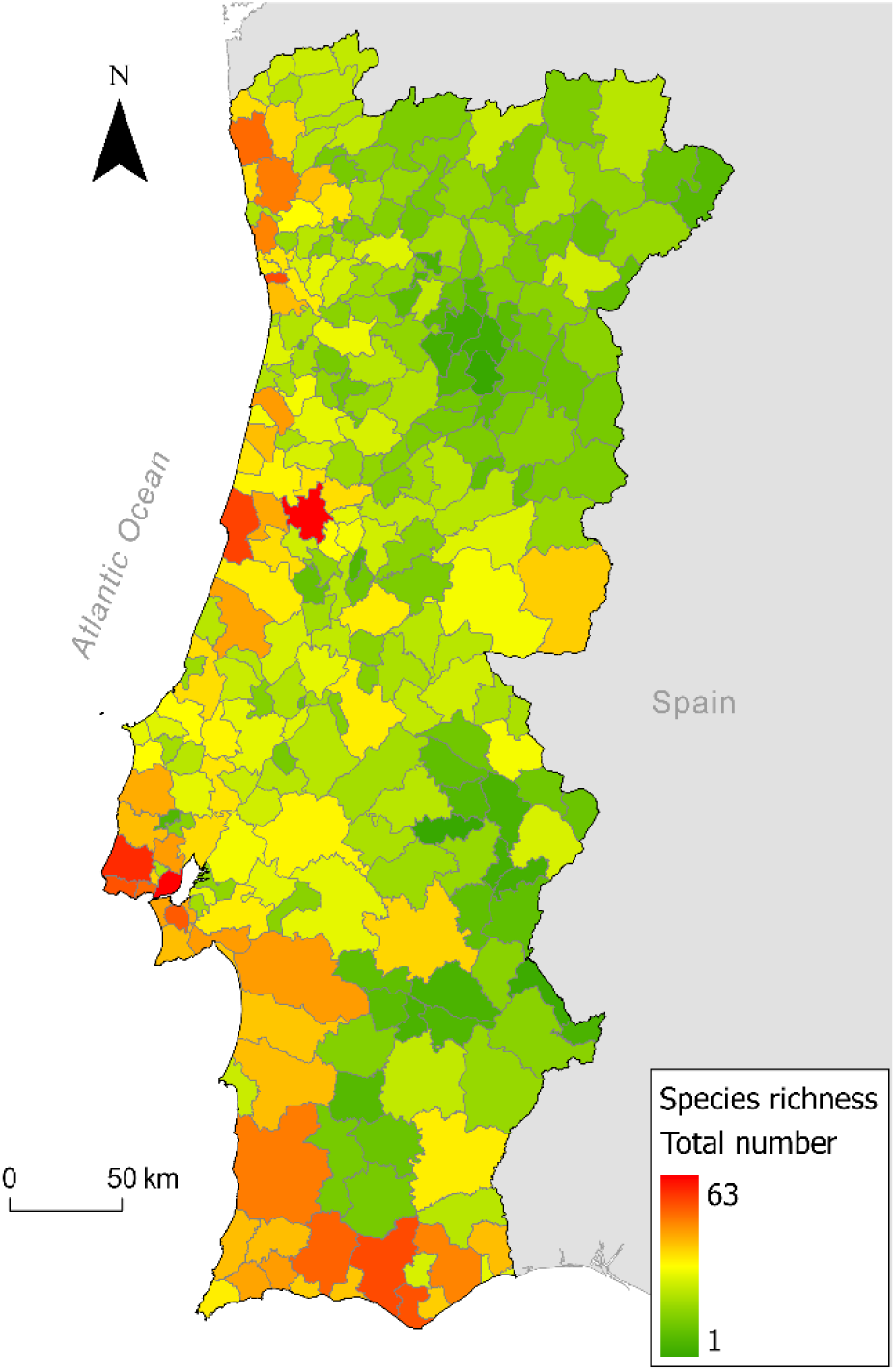
Richness of invasive alien plant species in mainland Portugal, corresponding to the total number of these species recorded in each municipality.

The GLS model, which aimed to identify the drivers of variation in species richness, achieved a pseudo-R^2^ value of 0.59, indicating good explanatory capacity. Four variables exhibited significant relationships with richness values (*p* < 0.05; Table 4, Suppl. material 3: Table S3.1). Specifically, species richness increased with larger municipality area, higher density of power lines, greater proximity to the coastline, and shorter travel time to cities.

**Table 4:** Coefficients and *p-value* results of each explanatory variable considered in the GLS model. Significant relationships (*p* < 0.05) with variations in invasive plants richness by municipality are shown in bold.

| Explanatory variable | Coefficient (SE) | *p*-value |
| --- | --- | --- |
| Daily mean air temperatures of the wettest quarter (°C) | -1.6352 (0.001) | 0.103 |
| Mean daily maximum air temperature of the warmest month (°C) | -1.1315 (0.029) | 0.259 |
| Ruggedness Index | 1.1009 (0.010) | 0.272 |
| **Minimum distance to coastline** | **-2.9901 (0.002)** | **0.003** |
| **Municipality area** | **8.4083 (0.006)** | **<.001** |
| Density of agricultural areas | 0.5164 (0.003) | 0.606 |
| Density of forest areas | 0.1772 (0.003) | 0.860 |
| Density of protected areas | 1.7760 (0.015) | 0.077 |
| Burnt area in the last 15 years | 0.1664 (0.002) | 0.868 |
| Soil pH | 0.6679 (1.176) | 0.505 |
| Density of local hydrographic network | 0.7588 (0.090) | 0.449 |
| Density of railways | 0.7601 (0.154) | 0.448 |
| **Density of power lines** | **2.0721 (0.231)** | **0.039** |
| **Travel time to cities (minutes)** | **-3.0433 (0.002)** | **0.003** |
| Sampling effort | 1.8438 (0.386) | 0.071 |
| Number of iNaturalist projects with IAP records | 0.8906 (0.189) | 0.374 |
| Number plant nurseries and aquarophilia stores | 1.7452 (0.045) | 0.082 |

### Discussion

Our assessment reveals that invasive alien plant species are now established throughout mainland Portugal, yet their patterns of distribution and occupancy are far from uniform. By integrating approximately 85,000 occurrence records from multiple sources, we identified the distribution of 96 invasive plant species across the study area. This represents a substantial increase in the knowledge about the distribution of invasive plant species richness in Portugal, once previous studies estimated the number of alien invasive plants present in Portugal (Almeida and Freitas 2000; Almeida and Freitas 2012; Morais et al. 2017) but did not aim to explore their distribution at the national scale. Other studies focused on national distribution but considered only a single species (e.g., González et al 2020).

#### Spatial patterns of species richness

While all municipalities host at least one invasive plant species, coastal and urbanized areas, particularly around Lisbon, Oporto, and the Algarve, emerge as concentrated hotspots of species richness. In contrast, inland and northeastern regions remain less heavily invaded. Our generalized least squares model identified proximity to the coastline and shorter travel times to urban centers as significant predictors of species richness, reinforcing the role of urbanization, human disturbance, and accessibility in driving invasions as favorable conditions for the introduction, establishment, and spread of alien species in many regions (e.g., Pyšek et al., 2010; Santos et al., 2011; Dawson et al., 2017; Pyšek et al., 2020; Dimitrakopoulos et al., 2022; Polce et al., 2023; Ivison et al., 2025). This reflects demographic and economic structure of Portugal, which is strongly coast-biased, with population and related activities mainly concentrated along the western and southern coastal regions, particularly in and around the Lisbon and Oporto metropolitan areas (Instituto Nacional de Estatística 2022). Additionally, some widespread invasive species in Portugal are known to have been brought initially to major urban centers as ornamental (e.g., *Ipomoea indica* or *Erigeron karvinskianus*; Marchante et al. 2014), while others were introduced specifically to coastal ecosystems in past decades, especially for dune fixation and erosion control (e.g., *Acacia longifolia* or *Carpobrotus edulis*; Marchante et al. 2014).

Species richness also correlated positively with both municipality size and the density of power-line corridors. The effect of area aligns with the classic species–area relationship, whereby larger spatial units tend to encompass a greater variety of habitats and a broader representation of the regional species pool, supporting more stable and abundant populations, and contributing to higher observed species richness (Matthews et al., 2021). On the other hand, linear infrastructures such as power lines may function as ecological corridors, especially considering that these areas are subject to vegetation cutting interventions, which cause disturbance and eliminate competition, factors known to facilitate the dispersal and establishment of invasive plant species (Dalu et al., 2023). Despite this, the role of such linear infrastructures in Portugal remains underexplored.

Despite climate being a known driver of alien species distributions (Dutra Silva et al. 2021), climatic variables were not significant predictors of invasive plant richness in this study. This may reflect the stronger influence of anthropogenic factors, namely human distribution and activities. Moreover, despite its relatively small size on global terms, mainland Portugal presents a relatively diverse climatic condition, shaped largely by its latitudinal gradient, with warmer dry Mediterranean-type climates in southern regions and colder temperate humid climate in northern coastal areas (Beck et al. 2023). We anticipated a significant relationship between soil pH and invasive species richness, given that calcareous and serpentinic soils—known for their high pH and challenging chemical profiles—tend to support fewer invasive plants. However, this expected effect may have been obscured in our analysis because municipality-level pH values represent averages across diverse soil types, thereby diluting the influence of localized extremes. Such soil heterogeneity can mask the inhibitory potential of highly alkaline or harsh serpentine soils on invasive species establishment.

Further insights emerged from the cluster analysis, which revealed four mainly distribution patterns, ranging from narrowly confined inland species to widely distributed taxa spanning both latitudinal and coastal-inland gradients. While most species conformed to expected patterns, notable exceptions emerged - e.g., *Cortaderia selloana*, was classified as widespread but is relatively uncommon in inland regions (González et al., 2020). This highlights how presence-only data at the municipal level may obscure finer-scale spatial variation.

These spatial and ecological patterns carry critical implications for monitoring and management. Species detected in only a few, et geographically dispersed municipalities — such as *Ludwigia grandiflora*—may indicate either early-stage invasions or underreporting. Differentiating between these scenarios is crucial, as early detection offers a window of opportunity for effective containment or eradication. Targeted field surveys and systematic monitoring should prioritize such cases to reduce uncertainty and support timely interventions.

Finally, a moderate positive association (R² = 0.42) between the number of municipalities a species occupies and its mean occupancy suggests that widespread species tend also to be locally dominant. This is evident for well-established invaders such as *Acacia dealbata*, *Arundo donax*, *Oxalis pes-caprae*, and *Phytolacca americana*, whose ecological impacts in mainland Portugal are well documented (Aguiar et al. 2013; Castro et al. 2016; Duarte et al. 2020; Marchante et al. 2023). In contrast, some species like *Acacia longifolia*, *Acacia saligna*, and *Carpobrotus edulis* show high local occupancy but limited national spread (mainly in coastal dune areas (Marchante et al. 2014)), suggesting environmentally constrained, yet intense, localized invasions.

#### Species richness as an indicator: opportunities and limitations

Our measure of species richness, derived from municipality-level presence data, provides a valuable and standardized overview of invasive plant distributions across mainland Portugal. While this approach does not capture local abundance—which could offer further insight into ecological dominance and impact—it remains a practical method for identifying general patterns of invasion. As with many large-scale biodiversity assessments, the underlying data are primarily sourced from opportunistic records, including citizen science contributions. Although these sources provide spatial coverage and volume of data, they can introduce spatial biases related to observer effort, site accessibility and species detectability (Tiago et al., 2017; Isaac et al., 2014; Westekemper et al., 2018). As a result, apparent absences in certain areas may reflect lower sampling effort rather than true species absence, particularly for less conspicuous or recently introduced taxa.

Moreover, species richness do not differentiate between species in terms of their ecological impact or management urgency. For example, the detection of *Fallopia japonica* in a new location may warrant immediate action due to its known severe ecological consequences (Aguilera et al 2010), whereas a new occurrence of *Erigeron karvinskianus* might pose a more localized and manageable concern. While our analysis adopts an equal-weight approach across species, it establishes a foundational framework that can be enhanced through species-specific risk assessments and prioritization strategies, which are vital for informed resource allocation and targeted management.

Despite limitations inherent in opportunistic and citizen science data (Robinson et al., 2017), citizen science data offer a cost-effective way to collect species records in greater numbers and in more locations. Moreover, using species richness as a pressure indicator is an helpful tool to identify areas where conservation and management efforts should be prioritized (Fleishman et al., 2006; Santos et al., 2011). However, while it is a useful tool for spotting potential invasion hotspots, it should not be interpreted as a direct measure of invasion severity or ecological impacts.

### Conclusion

This study provides insights about the spatial patterns and environmental and human drivers of invasive plant richness in mainland Portugal municipalities. Coastal urban areas show the highest species richness, largely driven by greater accessibility. This suggests that municipalities in these regions should prioritise invasive plant management due to their higher levels of impacts. At the same time, it is crucial that municipalities with fewer invasive species act early to prevent their spread, thereby increasing the effectiveness and sustainability of management efforts nationwide. Despite relying on the total number of invasive plants in each municipality, and not their abundance, the results offer a valuable baseline for identifying invasion hotspots and guiding management priorities. Continued monitoring and systematic data collection are essential to support effective prevention and control of plant invasions across the country.

## Supporting information

Results with all occurence data

Atlas of invasive plant species

Results without FloraOn data

## Acknowledgments

We would like to express our thanks to the many experts who have contributed by providing useful information on the occurrence records (Ascendi, Altri, Infraestruturas de Portugal, S.A., Liliana Duarte, Mónica Almeida, Sílvia Martins, MED-Mediterranean Institute for Agriculture, Environment and Development, and Botany Lab/Applied Ecology and Conservation Research Group of the University of Évora), the National Electricity Transmission Network shapefile, provided by David Almeida (Redes Energéticas Nacionais (REN)), and information on plant nurseries and aquarophilia stores data, collected by Ana Sofia Nunes (Nunes et al. unpublished).

## Additional information Conflict of interest

The authors have declared that no competing interests exist.

## Ethical statement

No ethical statement was reported.

## Author contributions

RF: Conceptualization, Data curation, Methodology, Writing – original draft, Writing – review and editing. AC: Methodology, Writing – original draft, Writing – review and editing. HM: Conceptualization, Supervision, Writing – review and editing. EM: Conceptualization, Supervision, Writing – review and editing. CC: Conceptualization, Supervision, Methodology, Writing – original draft, Writing – review and editing.

