## Supplementary material for "Distribution patterns of invasive alien plant species in mainland Portugal": Results with all occurence data

### Supplementary material 1

**Table S1.1.** Abbreviations and scientific names of each species considered in the database of occurrence records of invasive plants in mainland Portugal.

| Abbreviation | Species |
|---|---|
| AbM | *Abutilon theophrasti* Medik. |
| AcC | *Acacia cyclops* A.Cunn. ex G.Don |
| AcD | *Acacia dealbata* Link. |
| AcL | *Acacia longifolia* (Andrews) Willd. |
| AcM | *Acacia mearnsii* De Wild. |
| AcMl | *Acacia melanoxylon* R.Br. |
| AcR | *Acacia provincialis* A.Camus |
| AcP | *Acacia pycnantha* Benth. |
| AcS | *Acacia saligna* (Labill.) H.L.Wendl. |
| AN | *Acer negundo* L. |
| AgA | *Agave americana* L. |
| AgeA | *Ageratina adenophora* (Spreng.) R.M.King & H.Rob. |
| AiA | *Ailanthus altissima* (Mill.) Swingle |
| AlJ | *Albizia julibrissin* Durazz. |
| AlP | *Alternanthera philoxeroides* (Mart.) Griseb. |
| AmA | *Amaranthus albus* L. |
| AmB | *Amaranthus blitoides* S.Watson |
| AmBl | *Amaranthus blitum* L. ssp. *emarginatus* (Moq. ex Uline & Bray) Carretero, Muñoz Garmendia & Pedrol |
| ACa | *Amaranthus caudatus* L. |
| ACr | *Amaranthus cruentus* L. |
| AD | *Amaranthus deflexus* L. |
| AHy | *Amaranthus hybridus* L. |
| AHyp | *Amaranthus hypochondriacus* L. |
| AmM | *Amaranthus muricatus* (Moq.) Hieron. |
| AmP | *Amaranthus powellii* S.Watson |
| AmR | *Amaranthus retroflexus* L. |
| AV | *Amaranthus viridis* L. |
| ArS | *Araujia sericifera* Brot. |
| ArCa | *Arctotheca calendula* (L.) Levyns |
| ArD | *Arundo donax* L. |
| AsC | *Asclepias curassavica* L. |
| AsS | *Aster squamatus* Hieron. |
| AusS | *Austrocylindropuntia subulata* (Muehlenpf.) Backeb. |
| AzF | *Azolla filiculoides* Lam. |

| | |
|---|---|
| BH | *Baccharis halimifolia* L. |
| BS | *Baccharis spicata* (Lam.) Baill. |
| BA | *Bidens aurea* Sherff |
| BF | *Bidens frondosa* L. |
| BP | *Bidens pilosa* L. |
| CA | *Carpobrotus acinaciformis* (L.) L.Bolus |
| CE | *Carpobrotus edulis* (L.) N.E. Br. |
| PeV | *Cenchrus longisetus* M.C.Johnst. |
| PS | *Cenchrus setaceus* (Forssk.) Morrone |
| CoS | *Cortaderia selloana* (Schult. & Schult.f.) Asch. & Graebn. |
| CoC | *Cotula coronopifolia* L. |
| DS | *Datura stramonium* L. |
| EC | *Ehrharta calycina* Sm. |
| ElC | *Elodea canadensis* Michx. |
| ED | *Elodea densa* (Planch.) Casp. |
| EB | *Erigeron bonariensis* L. |
| ECa | *Erigeron canadensis* L. |
| EK | *Erigeron karvinskianus* DC. |
| CS | *Erigeron sumatrensis* Retz. |
| EP | *Eryngium pandanifolium* Cham. & Schltdl. |
| FB | *Fallopia baldschuanica* (Regel) Holub |
| GP | *Galinsoga parviflora* Cav. |
| GT | *Gleditsia triacanthos* L. |
| GF | *Gomphocarpus fruticosus* (L.) W.T.Aiton |
| GuT | *Gunnera tinctoria* (Molina) Mirb. |
| HD | *Hakea decurrens* R.Br. |
| HS | *Hakea salicifolia* (Vent.) B.L.Burtt |
| HG | *Hedychium gardnerianum* Sheph. ex Ker Gawl. |
| IpI | *Ipomoea indica* Merr. |
| SB | *Jacobaea maritima* subsp. Maritima |
| LM | *Lagarosiphon major* (Ridl.) Moss |
| LC | *Lantana camara* L. |
| LJ | *Lonicera japonica* Thunb. |
| LG | *Ludwigia grandiflora* (Michx.) Greuter & Burdet |
| LP | *Ludwigia peploides* (Kunth) P.H.Raven |
| MyA | *Myriophyllum aquaticum* (Vell.) Verdc. |
| NG | *Nicotiana glauca* Graham |
| NyM | *Nymphaea mexicana* Zucc. |
| OpE | *Opuntia elata* Link & Otto ex Salm-Dyck |
| OFI | *Opuntia ficus-indica* (L.) Mill. |
| OPC | *Oxalis pes-caprae* L. |
| OxP | *Oxalis purpurea* L. |
| PLo | *Paraserianthes lophantha* (Vent.) I.C.Nielsen |
| PP | *Paspalum distichum* L. |
| PVa | *Paspalum vaginatum* Sw. |
| PT | *Paulownia tomentosa* (Thunb.) Steud. |
| PyA | *Phytolacca americana* L. |

| | |
|---|---|
| PiS | *Pistia stratiotes* L. |
| PiU | *Pittosporum undulatum* Vent. |
| PoC | *Pontederia crassipes* Mart. |
| RJ | *Reynoutria japonica* Houtt. |
| RiC | *Ricinus communis* L. |
| RP | *Robinia pseudoacacia* L. |
| SM | *Salvinia molesta* D.Mitch. |
| SeI | *Senecio inaequidens* DC. |
| SoM | *Solanum mauritianum* Scop. |
| SoH | *Sorghum halepense* (L.) Pers. |
| SpD | *Sporobolus montevidensis* (Arechav.) P.M.Peterson & Saarela |
| TF | *Tradescantia fluminensis* Vell. |
| TrM | *Tropaeolum majus* L. |
| AcK | *Vachellia karroo* (Hayne) Banfi & Galasso (Synonym of *Acacia karroo* Hayne) |
| Wm | *Watsonia meriana* Mill. |

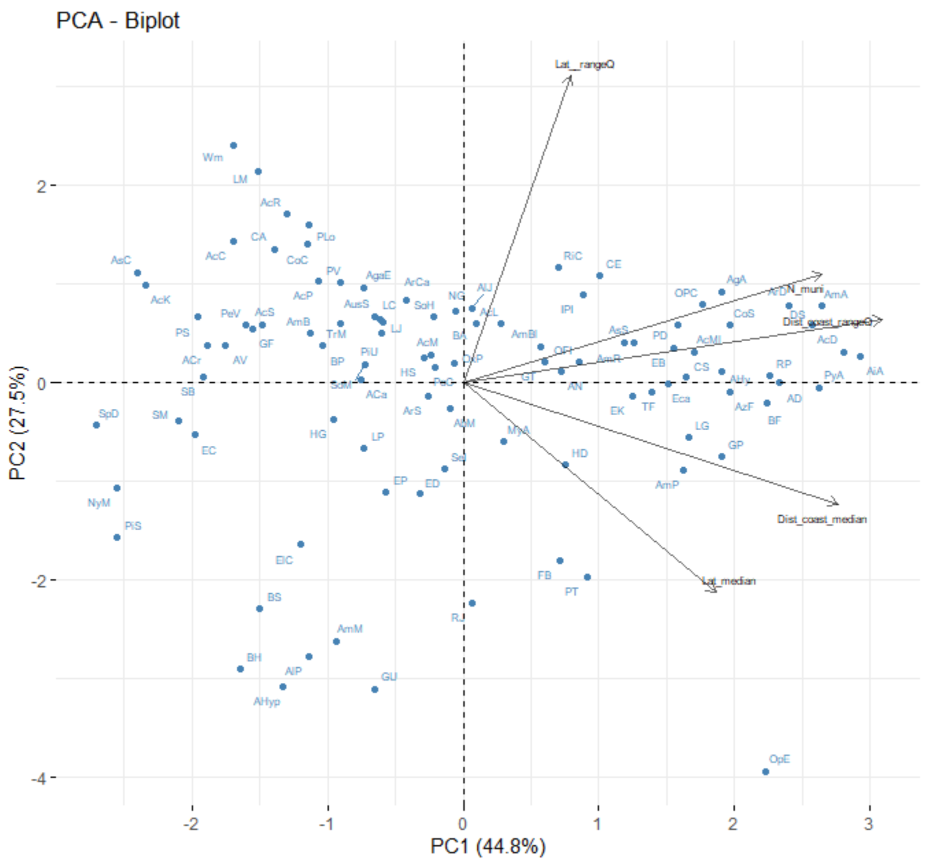


**Figure S1.1** Principal Component Analysis (PCA) biplot based on the complete invasive plants occurrence dataset. The first two Principal Components were computed in the biplot, explaining 72.3% of the variance. The PCA axes represent the total number of municipalities (N_muni), the median distances to the coast (Dist_coast_median), the interquartile range of distances to the coast (Dist_coast_rangeQ), the median latitudes (Lat_median), and the interquartile range of the latitudes (Lat_rangeQ).

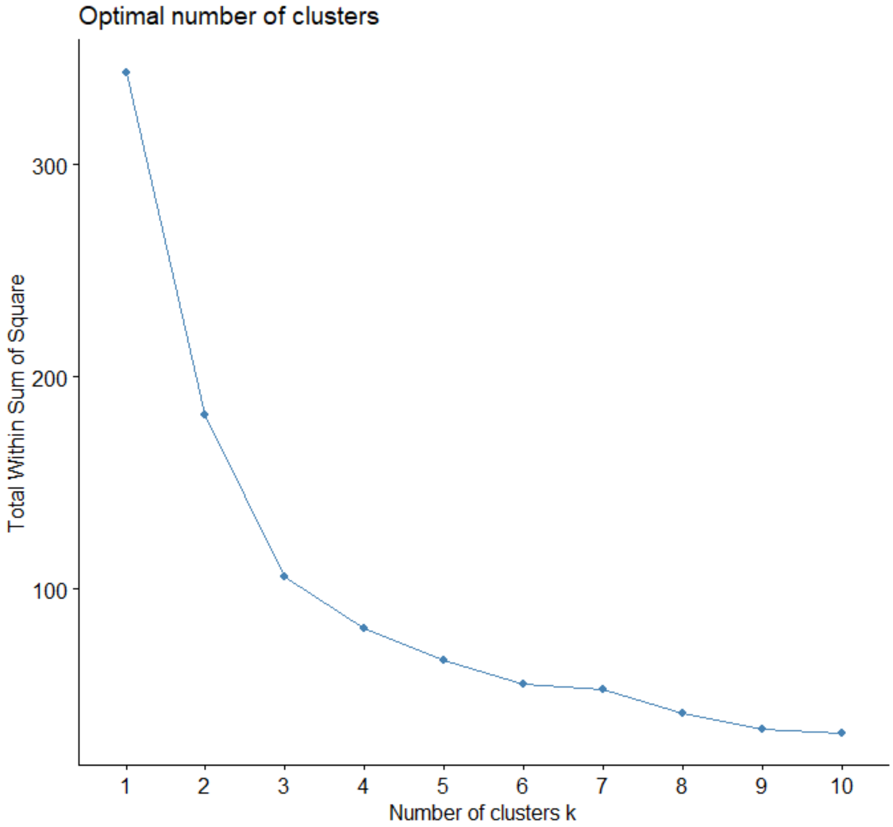


**Figure S1.2.** Optimal number of clusters based on the Elbow Method. Based on the method, the ideal number of clusters is four.

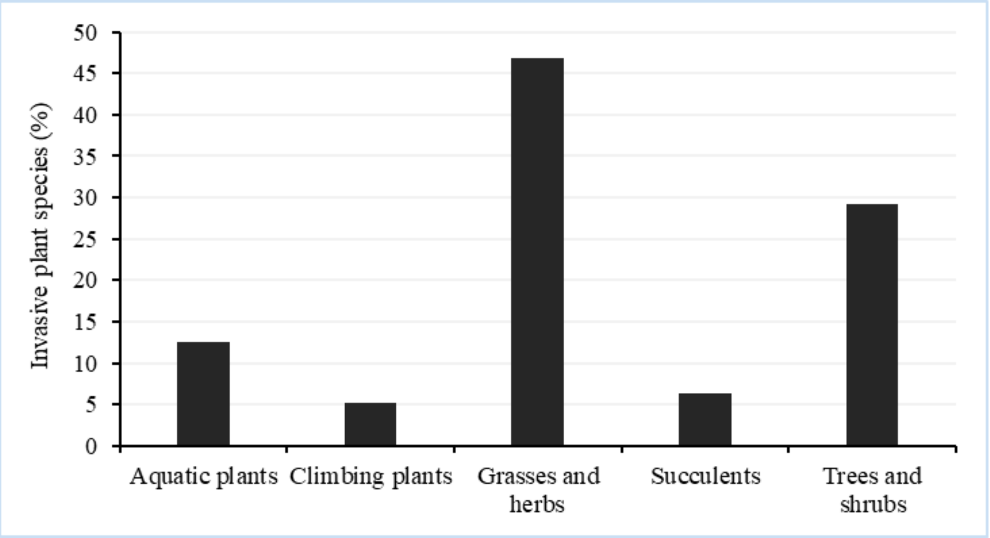


**Figure S1.3.** Growth form of the invasive alien plant species recorded in mainland Portugal.

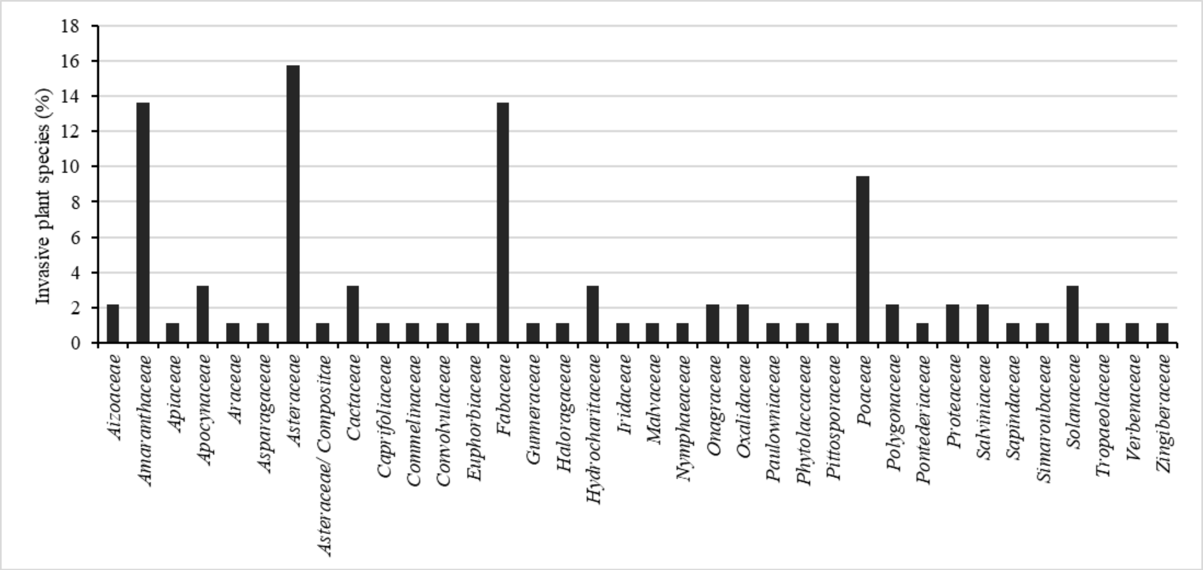


**Figure S1.4.** Families of the invasive alien plant species recorded as present in mainland Portugal.

**Table S1.2.** The 10 species of invasive plants with records in the fewest number of Portuguese municipalities.

| **Species** | **Number of municipalities with records of the species (frequency)** |
|---|---|
| *Baccharis spicata* | 4 |
| *Ludwigia grandiflora* | 4 |
| *Opuntia elata* | 3 |
| *Gunnera tinctoria* | 2 |
| *Nymphaea mexicana* | 2 |
| *Pistia stratiotes* | 2 |
| *Alternanthera philoxeroides* | 1 |
| *Amaranthus hypochondriacus* | 1 |
| *Amaranthus muricatus* | 1 |
| *Baccharis halimifolia* | 1 |

**Table S1.3.** Invasive plants present in mainland Portugal, sorted from the highest mean occupancy to the lowest. Mean occupancy was calculated as the percentage of 1×1 km grid cells occupied by each species within a municipality, averaged across all municipalities where the species was recorded.

| Species | Number of municipalities | Mean occupancy (%) |
|---|---|---|
| *Oxalis pes-caprae* | 213 | 7.53 |
| *Arundo donax* | 232 | 6.90 |
| *Cortaderia selloana* | 208 | 6.87 |
| *Arctotheca calendula* | 111 | 5.88 |
| *Acacia longifolia* | 131 | 5.58 |
| *Acacia dealbata* | 249 | 5.18 |
| *Carpobrotus edulis* | 136 | 5.13 |
| *Acacia saligna* | 52 | 4.92 |
| *Ricinus communis* | 137 | 4.61 |
| *Acacia melanoxylon* | 190 | 4.30 |
| *Pittosporum undulatum* | 85 | 4.20 |
| *Phytolacca americana* | 224 | 3.79 |
| *Ipomoea indica* | 164 | 3.67 |
| *Tropaeolum majus* | 81 | 3.54 |
| *Solanum mauritianum* | 14 | 3.09 |
| *Agave americana* | 187 | 2.93 |
| *Lantana camara* | 90 | 2.84 |
| *Tradescantia fluminensis* | 165 | 2.79 |
| *Erigeron bonariensis* | 159 | 2.75 |
| *Erigeron karvinskianus* | 149 | 2.59 |
| *Gomphocarpus fruticosus* | 31 | 2.53 |
| *Baccharis spicata* | 4 | 2.33 |
| *Reynoutria japonica* | 33 | 2.32 |
| *Ailanthus altissima* | 198 | 2.31 |
| *Datura stramonium* | 233 | 2.27 |
| *Araujia sericifera* | 64 | 2.21 |
| *Hakea salicifolia* | 27 | 2.14 |
| *Nicotiana glauca* | 57 | 2.06 |
| *Carpobrotus acinaciformis* | 48 | 2.01 |
| *Cenchrus setaceus* | 22 | 1.95 |
| *Ageratina adenophora* | 41 | 1.95 |
| *Baccharis halimifolia* | 1 | 1.89 |
| *Paraserianthes lophantha* | 54 | 1.88 |
| *Myriophyllum aquaticum* | 62 | 1.83 |
| *Eryngium pandanifolium* | 12 | 1.83 |
| *Cenchrus longisetus* | 22 | 1.79 |
| *Cotula coronopifolia* | 55 | 1.77 |
| *Acacia pycnantha* | 36 | 1.76 |
| *Oxalis purpurea* | 60 | 1.73 |
| *Pontederia crassipes* | 70 | 1.72 |
| *Robinia pseudoacacia* | 186 | 1.60 |
| *Bidens pilosa* | 37 | 1.60 |
| *Opuntia ficus-indica* | 99 | 1.57 |
| *Bidens aurea* | 110 | 1.55 |
| *Hakea decurrens* | 80 | 1.52 |
| *Watsonia meriana* | 8 | 1.44 |

| | | |
|---|---|---|
| *Acer negundo* | 91 | 1.43 |
| *Gleditsia triacanthos* | 54 | 1.43 |
| *Acacia provincialis* | 42 | 1.41 |
| *Lonicera japonica* | 69 | 1.38 |
| *Acacia mearnsii* | 69 | 1.35 |
| *Galinsoga parviflora* | 147 | 1.34 |
| *Vachellia karroo* | 12 | 1.31 |
| *Gunnera tinctoria* | 2 | 1.24 |
| *Amaranthus muricatus* | 1 | 1.20 |
| *Elodea densa* | 19 | 1.19 |
| *Aster squamatus* | 147 | 1.15 |
| *Bidens frondosa* | 178 | 1.15 |
| *Elodea canadensis* | 9 | 1.09 |
| *Paulownia tomentosa* | 16 | 1.05 |
| *Ludwigia peploides* | 8 | 1.04 |
| *Pistia stratiotes* | 2 | 1.04 |
| *Lagarosiphon major* | 12 | 1.01 |
| *Albizia julibrissin* | 32 | 1.01 |
| *Salvinia molesta* | 6 | 0.98 |
| *Acacia cyclops* | 10 | 0.98 |
| *Sporobolus montevidensis* | 10 | 0.96 |
| *Senecio inaequidens* | 7 | 0.89 |
| *Amaranthus blitoides* | 36 | 0.86 |
| *Austrocylindropuntia subulata* | 43 | 0.86 |
| *Amaranthus viridis* | 9 | 0.86 |
| *Hedychium gardnerianum* | 16 | 0.81 |
| *Amaranthus deflexus* | 108 | 0.81 |
| *Erigeron canadensis* | 99 | 0.78 |
| *Amaranthus caudatus* | 5 | 0.77 |
| *Sorghum halepense* | 29 | 0.71 |
| *Jacobaea maritima* | 15 | 0.70 |
| *Asclepias curassavica* | 10 | 0.69 |
| *Erigeron sumatrensis* | 105 | 0.67 |
| *Abutilon theophrasti* | 33 | 0.67 |
| *Amaranthus retroflexus* | 65 | 0.67 |
| *Ludwigia grandiflora* | 4 | 0.66 |
| *Paspalum distichum* | 123 | 0.66 |
| *Azolla filiculoides* | 56 | 0.63 |
| *Amaranthus hybridus* | 62 | 0.62 |
| *Amaranthus blitum* | 28 | 0.59 |
| *Amaranthus powellii* | 46 | 0.58 |
| *Paspalum vaginatum* | 14 | 0.55 |
| *Amaranthus albus* | 44 | 0.50 |
| *Ehrharta calycina* | 7 | 0.49 |
| *Fallopia baldschuanica* | 5 | 0.49 |
| *Amaranthus cruentus* | 7 | 0.46 |
| *Opuntia elata* | 3 | 0.42 |
| *Nymphaea mexicana* | 2 | 0.40 |
| *Alternanthera philoxeroides* | 1 | 0.31 |
| *Amaranthus hypochondriacus* | 1 | 0.27 |

**Table S1.4.** Attributed cluster to each invasive plant present in mainland Portugal and the variables that characterize their distribution, based on the municipalities where they are recorded: the number of municipalities, median and interquartile ranges of distances to the coast (km), and median and interquartile ranges of latitudes (decimal degrees). Cluster 1 encompasses the species primarily located along the coast. Cluster 2 mostly comprises species with narrow ranges. Cluster 3 includes the moderately widespread species. Cluster 4 is the widespread species.

| Abbreviation | Species | Cluster | Number of municipalities | Median distances to the coast (km) | Interquartile range of distances to the coast (km) | Median latitudes (decimal degrees) | Interquartile range of the latitudes (decimal degrees) |
|---|---|---|---|---|---|---|---|
| AcC | *Acacia cyclops* | 1 | 10 | 3.67 | 14.80 | 38.68 | 2.75 |
| AcP | *Acacia pycnantha* | 1 | 36 | 11.95 | 42.41 | 38.01 | 1.88 |
| AcR | *Acacia retinodes* | 1 | 42 | 7.53 | 14.60 | 38.71 | 2.98 |
| AcS | *Acacia saligna* | 1 | 52 | 8.34 | 13.72 | 38.63 | 1.80 |
| AgaE | *Ageratina adenophora* | 1 | 41 | 16.31 | 25.45 | 39.16 | 2.56 |
| AmB | *Amaranthus blitoides* | 1 | 36 | 16.81 | 23.03 | 38.66 | 1.88 |
| ACr | *Amaranthus cruentus* | 1 | 7 | 9.06 | 13.26 | 38.46 | 1.69 |
| AV | *Amaranthus viridis* | 1 | 9 | 10.89 | 11.54 | 38.66 | 1.82 |
| AsC | *Asclepias curassavica* | 1 | 10 | 5.50 | 11.94 | 37.27 | 1.76 |
| BP | *Bidens pilosa* | 1 | 37 | 16.31 | 22.18 | 38.96 | 1.89 |
| CA | *Carpobrotus acinaciformis* | 1 | 49 | 7.09 | 13.21 | 38.65 | 2.57 |
| PeV | *Cenchrus longisetus* | 1 | 22 | 7.94 | 14.35 | 38.80 | 1.99 |
| PS | *Cenchrus setaceus* | 1 | 22 | 7.30 | 14.26 | 38.05 | 1.67 |
| CoC | *Cotula coronopifolia* | 1 | 55 | 7.37 | 12.58 | 39.09 | 2.84 |
| EC | *Ehrharta calycina* | 1 | 7 | 11.52 | 9.28 | 38.66 | 0.95 |
| GF | *Gomphocarpu s fruticosus* | 1 | 31 | 7.75 | 15.43 | 38.76 | 1.89 |
| SB | *Jacobaea maritima* | 1 | 15 | 6.07 | 5.82 | 38.96 | 1.59 |
| LM | *Lagarosiphon major* | 1 | 12 | 16.29 | 22.79 | 37.49 | 3.00 |

| | | | | | | | |
|---|---|---|---|---|---|---|---|
| NyM | *Nymphaea mexicana* | 1 | 2 | 22.14 | 0.00 | 37.36 | 0.00 |
| PLo | *Paraserianthes lophantha* | 1 | 54 | 8.38 | 15.91 | 38.80 | 2.88 |
| PV | *Paspalum vaginatum* | 1 | 14 | 11.97 | 13.01 | 40.03 | 3.16 |
| SM | *Salvinia molesta* | 1 | 6 | 5.83 | 8.05 | 38.80 | 1.08 |
| SpD | *Sporobolus montevidensis* | 1 | 10 | 6.29 | 12.07 | 37.21 | 0.18 |
| TrM | *Tropaeolum majus* | 1 | 81 | 9.27 | 16.58 | 39.28 | 2.02 |
| AcK | *Vachellia karroo* | 1 | 12 | 6.29 | 14.84 | 37.25 | 1.60 |
| Wm | *Watsonia meriana* | 1 | 8 | 14.99 | 14.52 | 37.49 | 3.30 |
| AlP | *Alternanthera philoxeroides* | 2 | 1 | 36.12 | 0.00 | 40.22 | 0.00 |
| AHyp | *Amaranthus hypochondriac us* | 2 | 1 | 13.86 | 0.00 | 41.54 | 0.00 |
| AmM | *Amaranthus muricatus* | 2 | 1 | 53.20 | 0.00 | 39.38 | 0.00 |
| BH | *Baccharis halimifolia* | 2 | 1 | 2.90 | 0.00 | 41.55 | 0.00 |
| BS | *Baccharis spicata* | 2 | 4 | 4.25 | 5.13 | 41.23 | 0.43 |
| ElC | *Elodea canadensis* | 2 | 9 | 17.06 | 15.29 | 40.16 | 0.66 |
| ED | *Elodea densa* | 2 | 19 | 18.95 | 26.21 | 41.09 | 1.59 |
| EP | *Eryngium pandanifolium* | 2 | 12 | 22.22 | 32.23 | 40.12 | 1.14 |
| FB | *Fallopia baldschuanica* | 2 | 5 | 32.33 | 59.13 | 41.40 | 1.14 |
| GU | *Gunnera tinctoria* | 2 | 2 | 49.88 | 0.00 | 40.44 | 0.00 |
| OpE | *Opuntia elata* | 2 | 3 | 111.34 | 32.72 | 41.23 | 0.41 |
| PT | *Paulownia tomentosa* | 2 | 16 | 45.78 | 51.61 | 41.13 | 1.06 |
| PiS | *Pistia stratiotes* | 2 | 2 | 2.68 | 0.00 | 38.98 | 0.00 |
| RJ | *Reynoutria japonica* | 2 | 33 | 27.24 | 34.63 | 41.24 | 0.59 |

| | | | | | | | |
|---|---|---|---|---|---|---|---|
| AbM | *Abutilon theophrasti* | 3 | 33 | 27.24 | 43.52 | 39.63 | 1.68 |
| AcL | *Acacia longifolia* | 3 | 131 | 18.16 | 32.89 | 39.78 | 2.15 |
| AcM | *Acacia mearnsii* | 3 | 69 | 16.33 | 44.98 | 39.25 | 1.70 |
| AN | *Acer negundo* | 3 | 91 | 31.22 | 48.25 | 39.91 | 2.00 |
| AlJ | *Albizia julibrissin* | 3 | 32 | 20.35 | 47.15 | 39.87 | 2.69 |
| AmBl | *Amaranthus blitum* | 3 | 28 | 38.30 | 64.59 | 39.12 | 2.13 |
| ACa | *Amaranthus caudatus* | 3 | 5 | 9.06 | 20.10 | 40.82 | 2.53 |
| ArS | *Araujia sericifera* | 3 | 64 | 13.76 | 21.25 | 40.74 | 2.16 |
| ArCa | *Arctotheca calendula* | 3 | 111 | 13.63 | 19.81 | 39.34 | 2.23 |
| AusS | *Austrocylindro puntia subulata* | 3 | 43 | 9.06 | 54.20 | 38.66 | 1.71 |
| BA | *Bidens aurea* | 3 | 110 | 17.38 | 39.92 | 39.42 | 1.98 |
| GT | *Gleditsia triacanthos* | 3 | 54 | 30.48 | 67.99 | 39.29 | 1.81 |
| HD | *Hakea decurrens* | 3 | 80 | 40.04 | 41.39 | 40.24 | 1.46 |
| HS | *Hakea salicifolia* | 3 | 27 | 19.72 | 40.95 | 39.72 | 2.16 |
| HG | *Hedychium gardnerianum* | 3 | 16 | 11.56 | 26.45 | 39.92 | 1.62 |
| IPI | *Ipomoea indica* | 3 | 164 | 21.94 | 45.23 | 39.67 | 2.24 |
| LC | *Lantana camara* | 3 | 90 | 11.20 | 28.76 | 39.08 | 1.89 |
| LJ | *Lonicera japonica* | 3 | 69 | 11.52 | 30.57 | 39.34 | 2.07 |
| LP | *Ludwigia peploides* | 3 | 8 | 20.40 | 22.82 | 40.18 | 1.67 |
| MyA | *Myriophyllum aquaticum* | 3 | 62 | 25.97 | 39.64 | 40.47 | 1.66 |
| NG | *Nicotiana glauca* | 3 | 57 | 17.43 | 66.08 | 38.59 | 1.75 |
| OFI | *Opuntia ficus-indica* | 3 | 99 | 28.97 | 73.28 | 39.07 | 1.45 |

| | | | | | | | |
|---|---|---|---|---|---|---|---|
| OxP | *Oxalis purpurea* | 3 | 60 | 19.41 | 33.18 | 40.12 | 2.21 |
| PiU | *Pittosporum undulatum* | 3 | 85 | 15.29 | 25.31 | 39.09 | 1.87 |
| PoC | *Pontederia crassipes* | 3 | 70 | 17.20 | 30.99 | 39.94 | 2.02 |
| RiC | *Ricinus communis* | 3 | 137 | 21.46 | 48.35 | 39.43 | 2.48 |
| SeI | *Senecio inaequidens* | 3 | 7 | 13.86 | 39.11 | 41.34 | 1.85 |
| SoM | *Solanum mauritianum* | 3 | 14 | 8.08 | 27.64 | 40.40 | 2.37 |
| SoH | *Sorghum halepense* | 3 | 29 | 18.81 | 49.19 | 39.35 | 2.31 |
| AcD | *Acacia dealbata* | 4 | 249 | 45.62 | 64.50 | 40.35 | 1.96 |
| AcMl | *Acacia melanoxylon* | 4 | 189 | 31.22 | 53.12 | 40.35 | 2.02 |
| AgA | *Agave americana* | 4 | 187 | 35.15 | 67.65 | 39.60 | 2.24 |
| AiA | *Ailanthus altissima* | 4 | 198 | 50.30 | 80.01 | 40.33 | 2.09 |
| AmA | *Amaranthus albus* | 4 | 44 | 79.04 | 96.04 | 38.92 | 2.88 |
| AD | *Amaranthus deflexus* | 4 | 108 | 53.28 | 82.07 | 40.14 | 2.12 |
| AHy | *Amaranthus hybridus* | 4 | 62 | 44.62 | 86.96 | 40.22 | 2.29 |
| AmP | *Amaranthus powellii* | 4 | 46 | 45.63 | 84.53 | 40.29 | 1.41 |
| AmR | *Amaranthus retroflexus* | 4 | 65 | 25.47 | 87.93 | 39.80 | 2.02 |
| ArD | *Arundo donax* | 4 | 232 | 39.00 | 65.90 | 39.91 | 2.15 |
| AsS | *Aster squamatus* | 4 | 147 | 30.40 | 59.19 | 39.72 | 1.91 |
| AzF | *Azolla filiculoides* | 4 | 56 | 54.74 | 87.50 | 39.78 | 2.04 |
| BF | *Bidens frondosa* | 4 | 179 | 46.74 | 61.38 | 40.41 | 1.78 |
| CE | *Carpobrotus edulis* | 4 | 136 | 19.41 | 62.24 | 39.66 | 2.40 |
| CoS | *Cortaderia selloana* | 4 | 208 | 31.37 | 59.38 | 40.22 | 2.12 |

| | | | | | | | |
|---|---|---|---|---|---|---|---|
| DS | *Datura stramonium* | 4 | 233 | 38.85 | 69.17 | 40.22 | 2.09 |
| EB | *Erigeron bonariensis* | 4 | 159 | 35.15 | 58.12 | 39.91 | 1.99 |
| Eca | *Erigeron canadensis* | 4 | 99 | 39.78 | 70.80 | 39.94 | 1.88 |
| EK | *Erigeron karvinskianus* | 4 | 150 | 32.44 | 45.77 | 40.35 | 1.79 |
| CS | *Erigeron sumatrensis* | 4 | 105 | 42.15 | 75.81 | 39.72 | 1.82 |
| GP | *Galinsoga parviflora* | 4 | 148 | 41.66 | 60.63 | 40.75 | 1.46 |
| LG | *Ludwigia grandiflora* | 4 | 4 | 39.54 | 116.75 | 39.81 | 1.38 |
| OPC | *Oxalis pes-caprae* | 4 | 213 | 32.54 | 58.87 | 39.57 | 2.00 |
| PD | *Paspalum distichum* | 4 | 123 | 39.19 | 72.08 | 39.48 | 2.14 |
| PyA | *Phytolacca americana* | 4 | 225 | 44.85 | 64.79 | 40.50 | 1.76 |
| RP | *Robinia pseudoacacia* | 4 | 186 | 42.09 | 65.70 | 40.38 | 1.91 |
| TF | *Tradescantia fluminensis* | 4 | 165 | 31.52 | 48.98 | 40.35 | 1.74 |
