## Supplementary material for "Distribution patterns of invasive alien plant species in mainland Portugal": Atlas of invasive plant species

**Supplementary material 2**

**Atlas of invasive plant species**

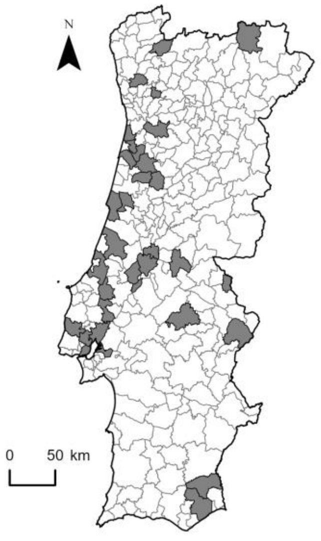


*Abutilon theophrasti*

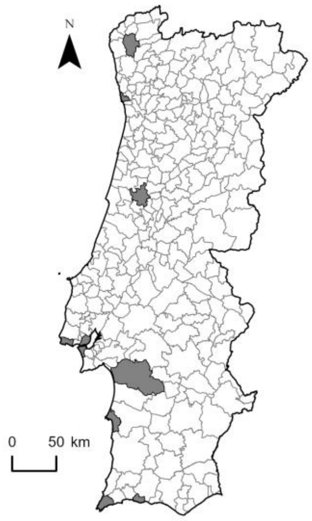


*Acacia cyclops*

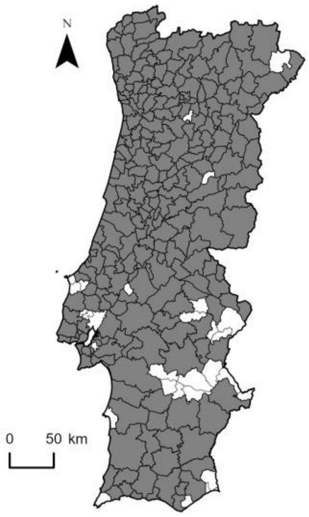


*Acacia dealbata*

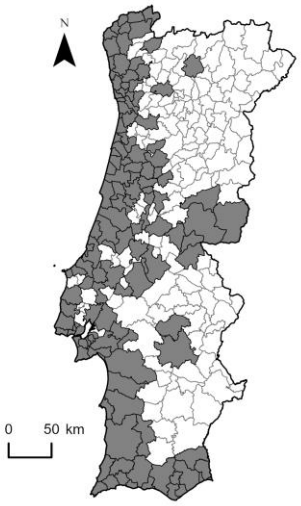


*Acacia longifolia*

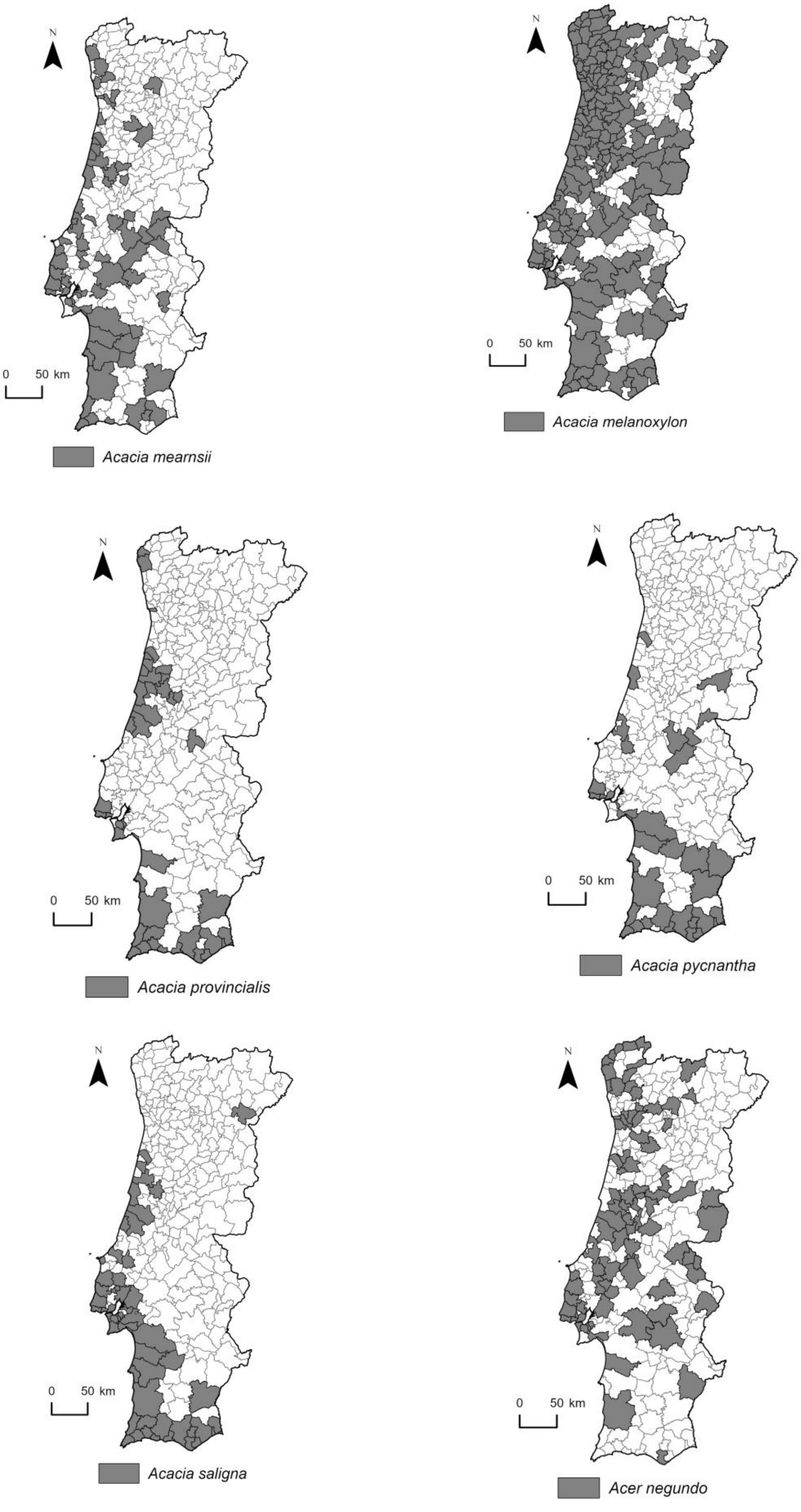
N
0 50 km
Acacia mearnsii
N
0 50 km
Acacia melanoxylon
N
0 50 km
Acacia provincialis
N
0 50 km
Acacia pycnantha
N
0 50 km
Acacia saligna
N
0 50 km
Acer negundo

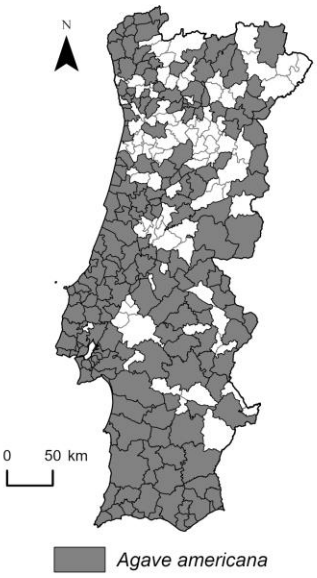
N
0
50 km
Agave americana

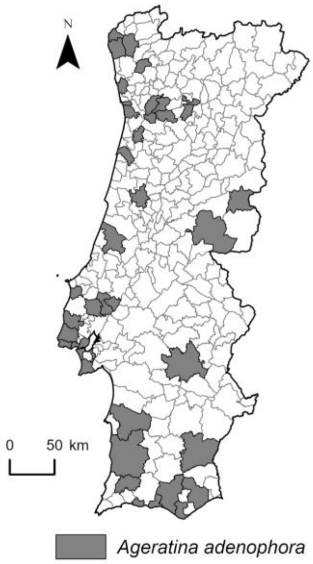
N
0
50 km
Ageratina adenophora

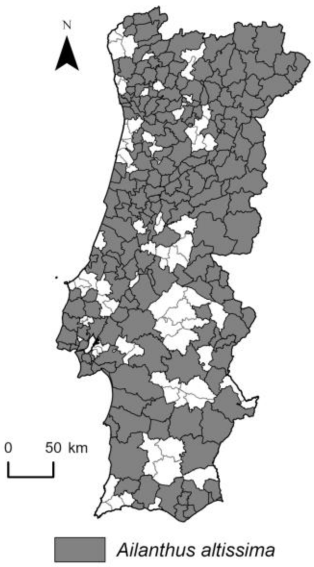
N
0
50 km
Ailanthus altissima

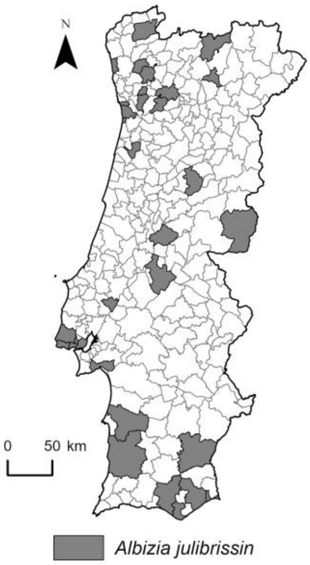
N
0
50 km
Albizia julibrissin

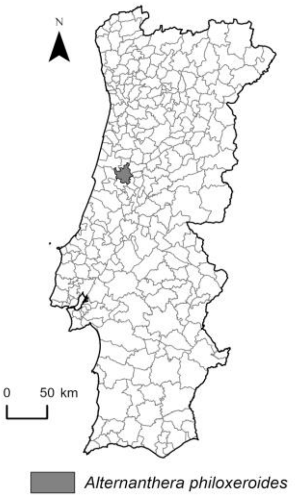
N
0
50 km
Alternanthera philoxeroides

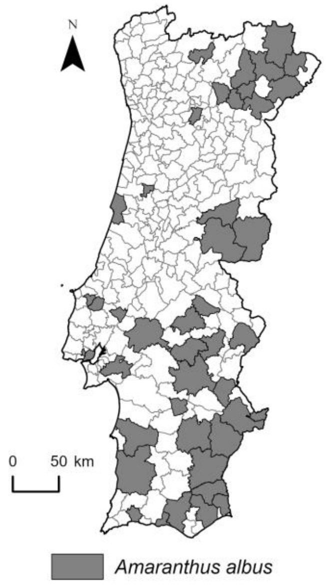
N
0 50 km
*Amaranthus albus*

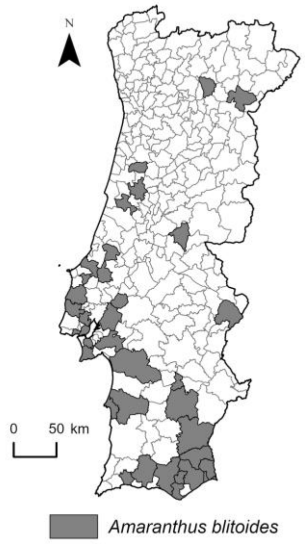
N
0 50 km
*Amaranthus blitoides*

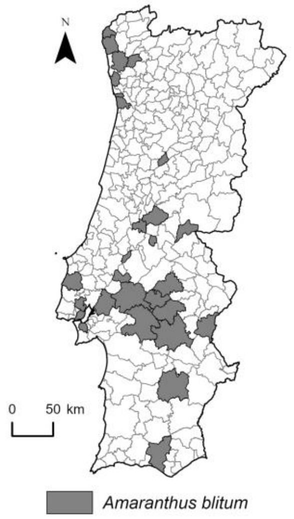
N
0 50 km
*Amaranthus blitum*

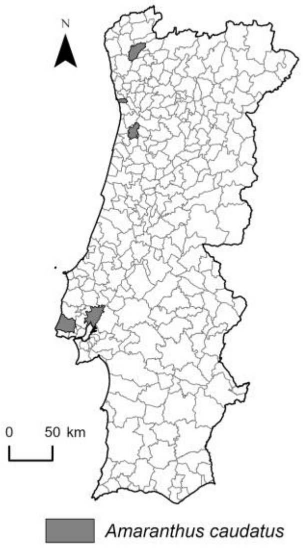
N
0 50 km
*Amaranthus caudatus*

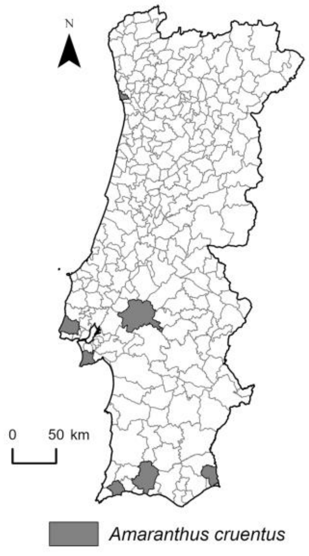
N
0 50 km
*Amaranthus cruentus*

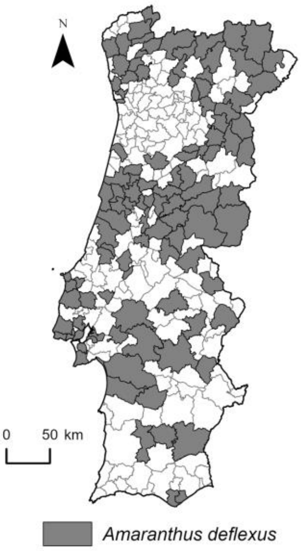
N
0
50 km
Amaranthus deflexus

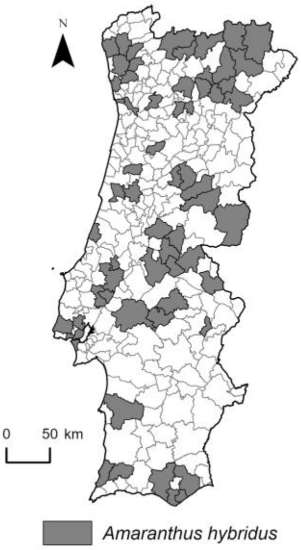
N
0
50 km
Amaranthus hybridus

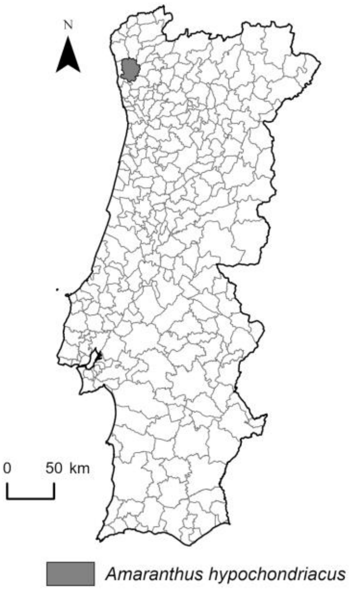
N
0
50 km
Amaranthus hypochondriacus

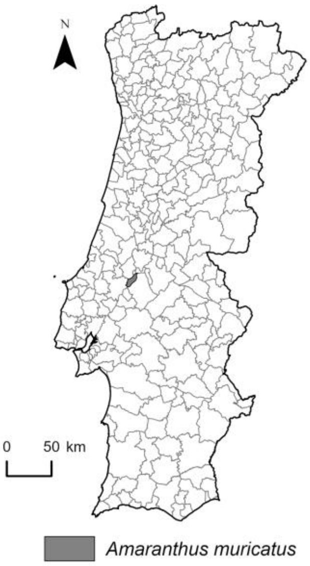
N
0
50 km
Amaranthus muricatus

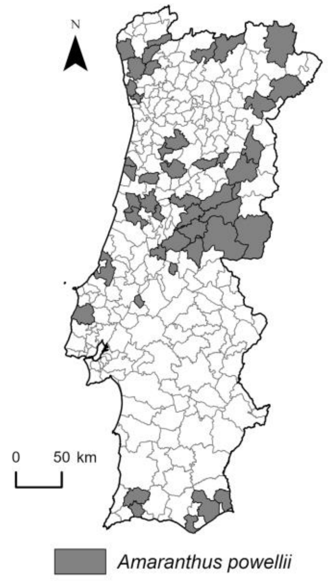
N
0 50 km
Amaranthus powellii

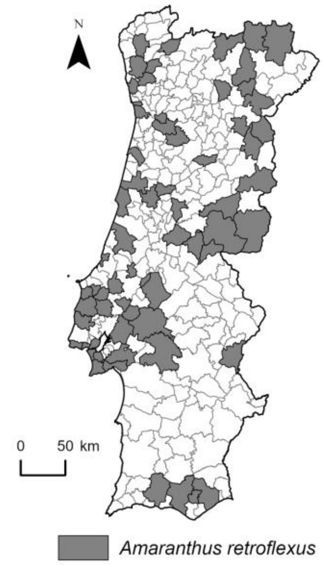
N
0 50 km
Amaranthus retroflexus

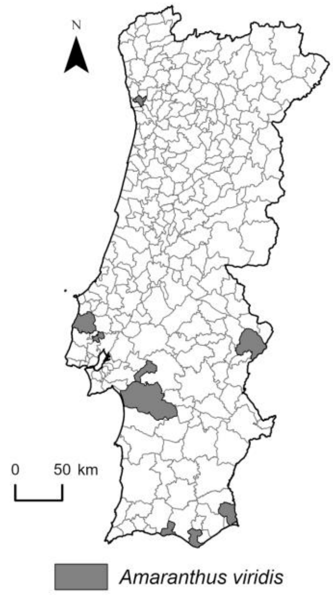
N
0 50 km
Amaranthus viridis

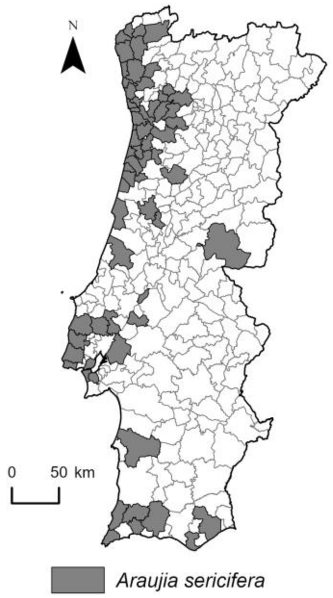
N
0 50 km
Araujia sericifera

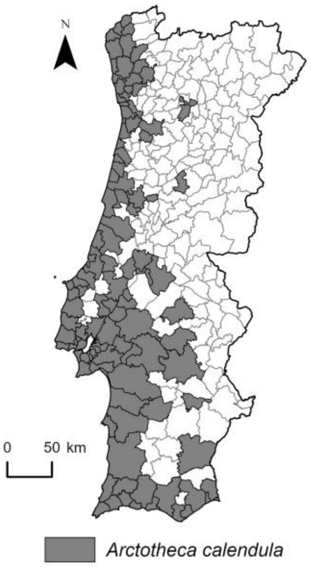
N
0 50 km
Arctotheca calendula

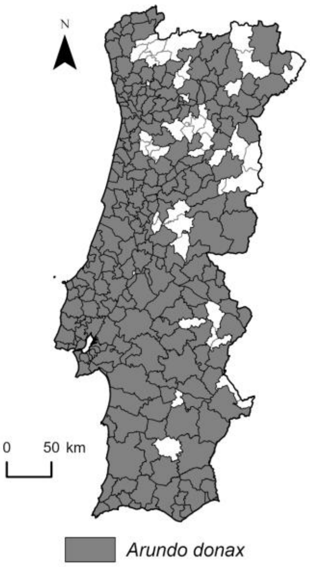
N
0 50 km
Arundo donax

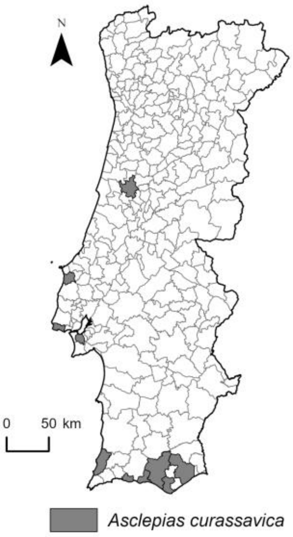
N
0 50 km
Asclepias curassavica

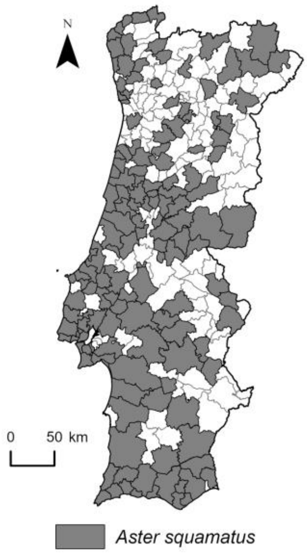
N
0 50 km
Aster squamatus

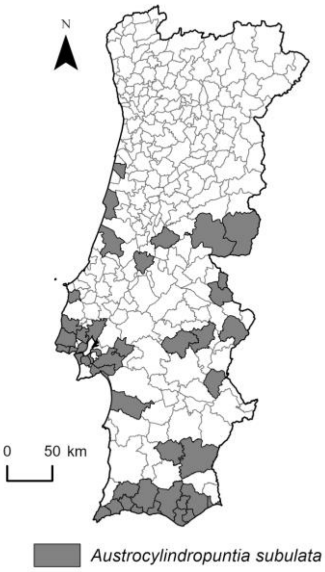
N
0 50 km
Austrocylindropuntia subulata

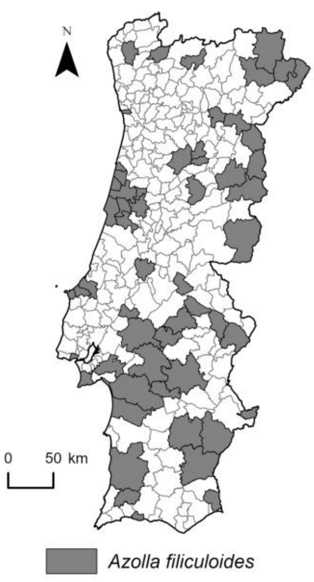
N
0 50 km
Azolla filiculoides

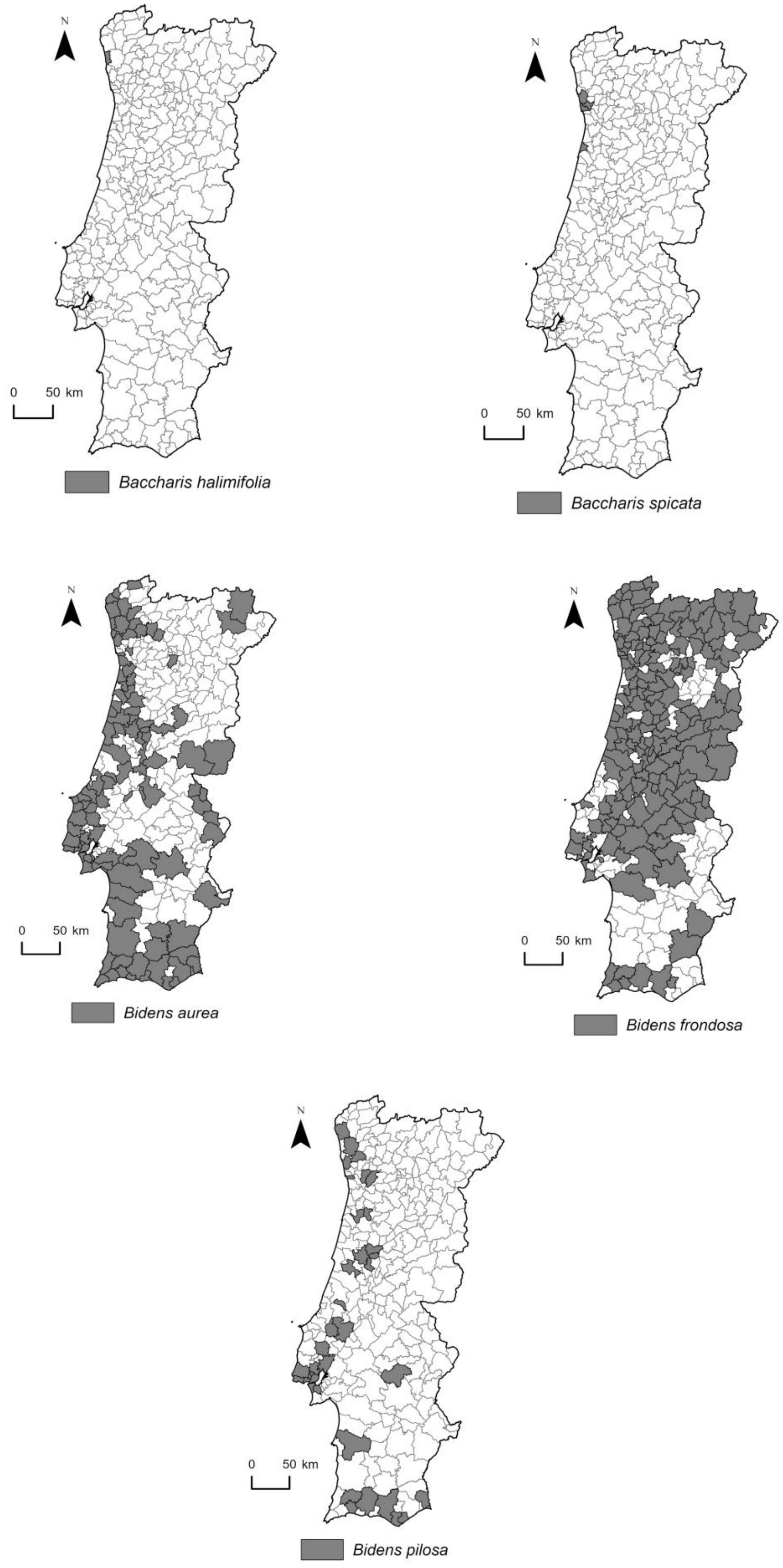
N
0 50 km
Baccharis halimifolia
N
0 50 km
Baccharis spicata
N
0 50 km
Bidens aurea
N
0 50 km
Bidens frondosa
N
0 50 km
Bidens pilosa

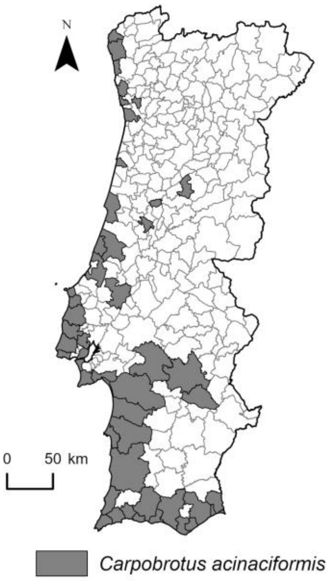
N
0
50 km
Carpobrotus acinaciformis

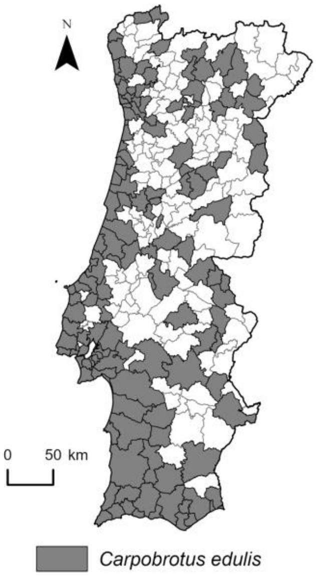
N
0
50 km
Carpobrotus edulis

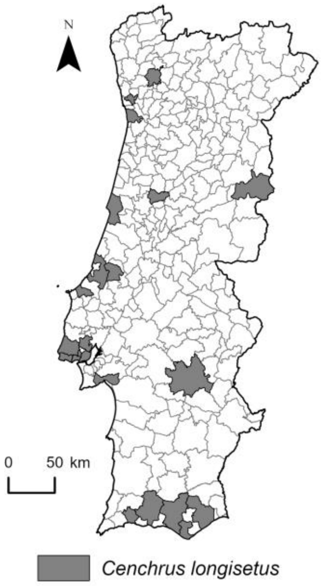
N
0
50 km
Cenchrus longisetus

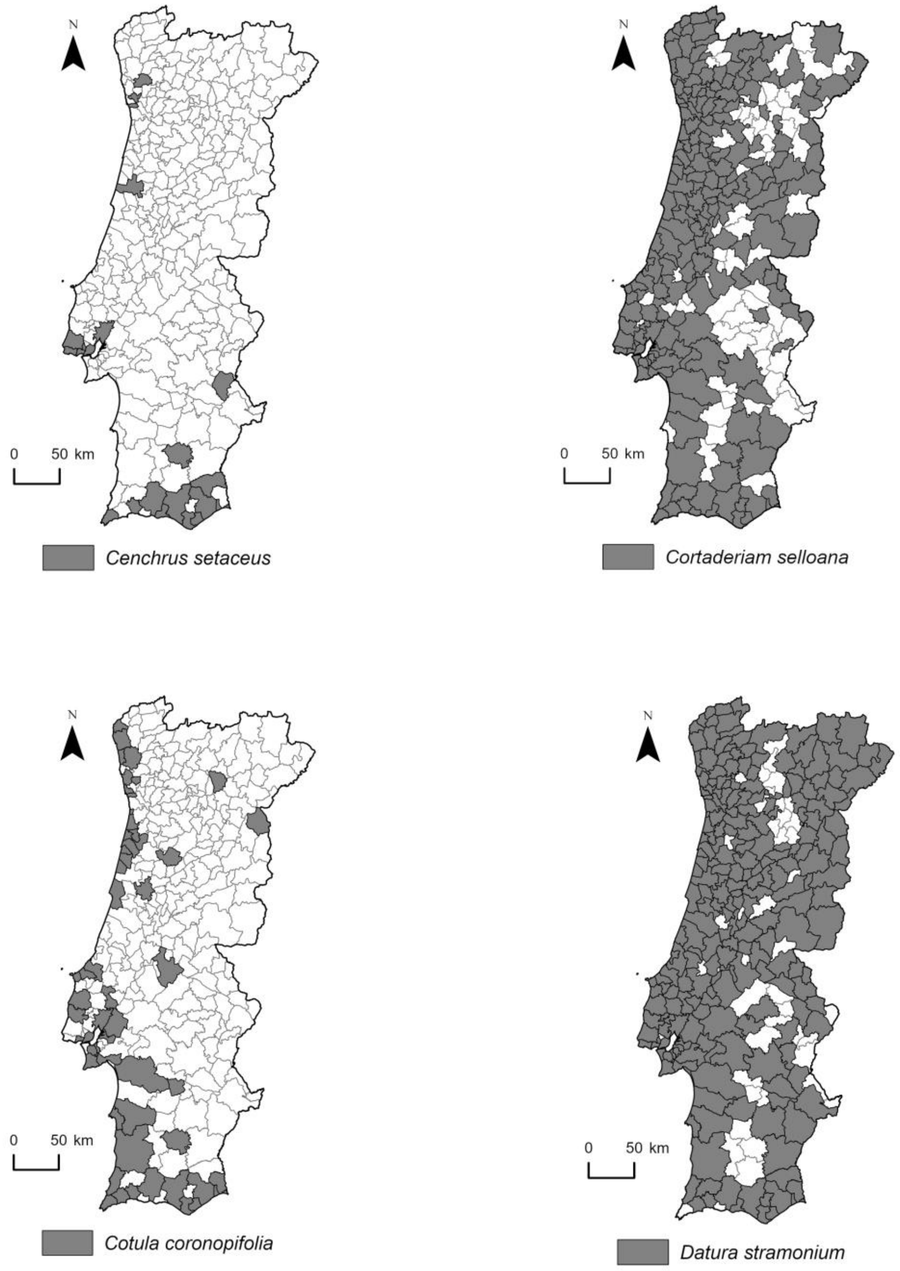
N
0
50 km
Cenchrus setaceus
N
0
50 km
Cortaderiam selloana
N
0
50 km
Cotula coronopifolia
N
0
50 km
Datura stramonium

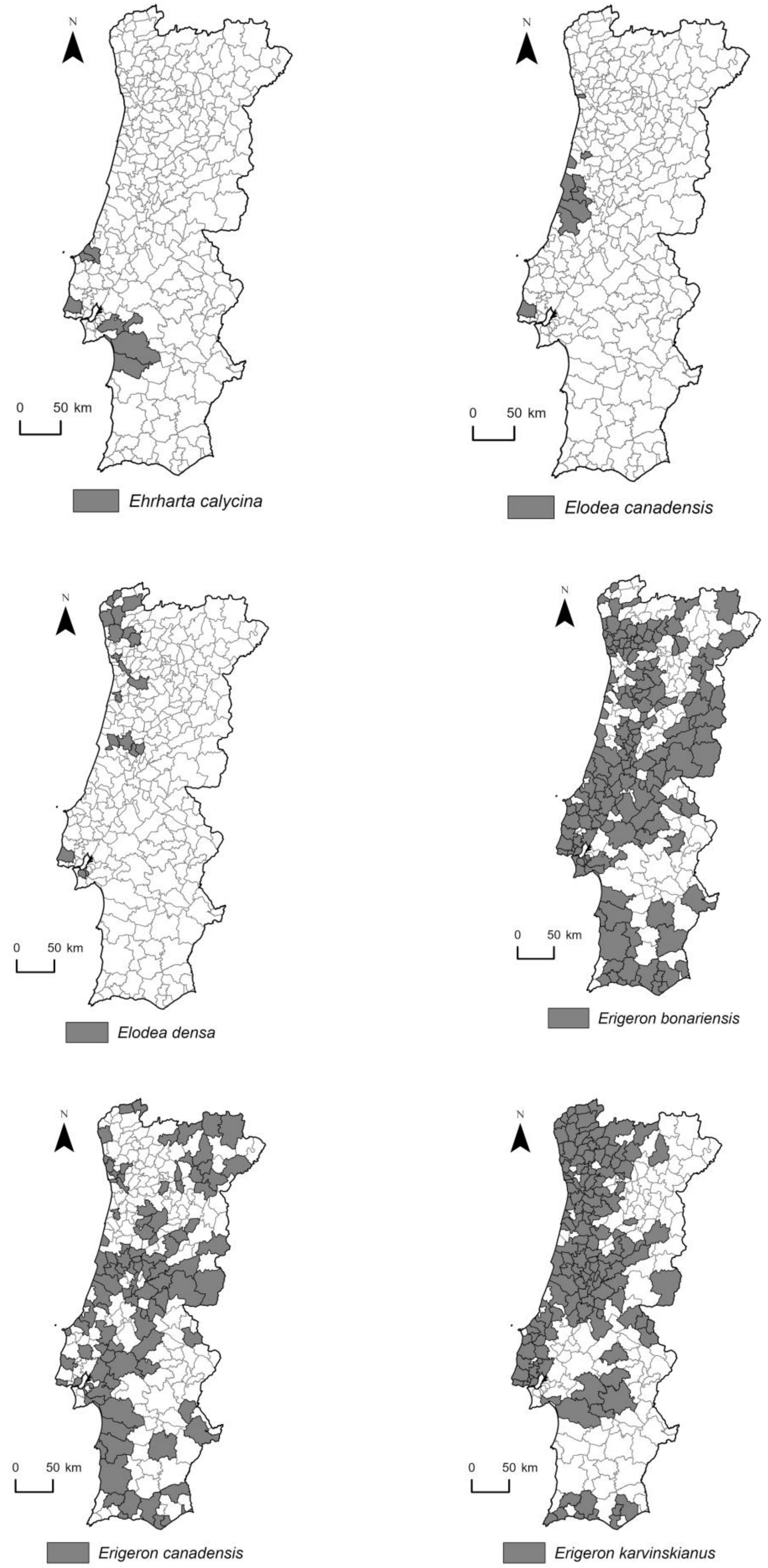
N
0 50 km
Ehrharta calycina
N
0 50 km
Elodea canadensis
N
0 50 km
Elodea densa
N
0 50 km
Erigeron bonariensis
N
0 50 km
Erigeron canadensis
N
0 50 km
Erigeron karvinskianus

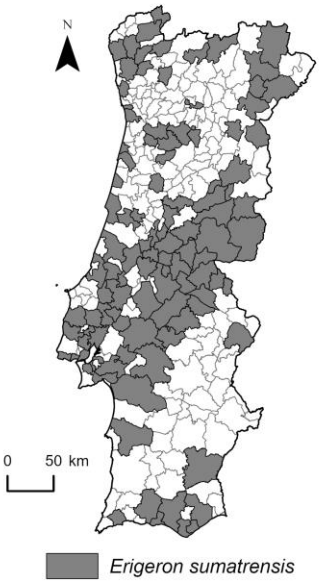
N
0
50 km
Erigeron sumatrensis

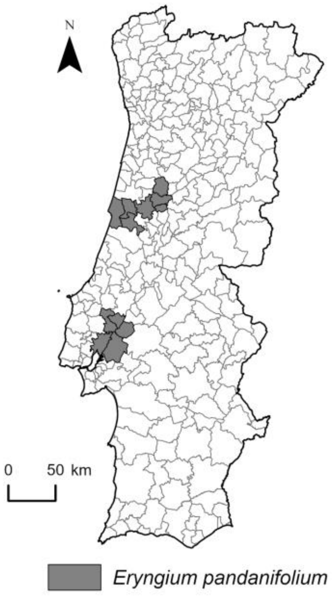
N
0
50 km
Eryngium pandanifolium

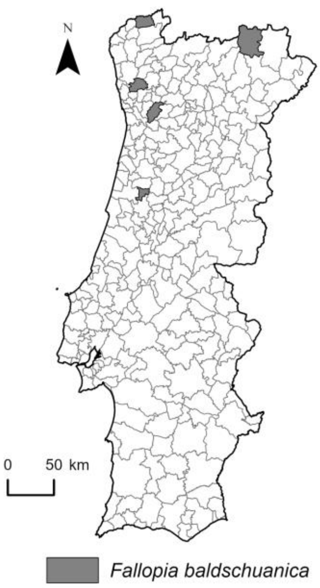
N
0
50 km
Fallopia baldschuanica

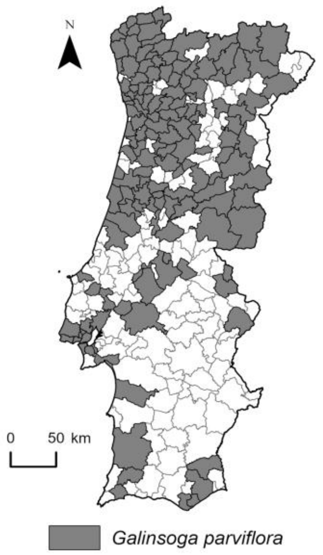
N
0
50 km
Galinsoga parviflora

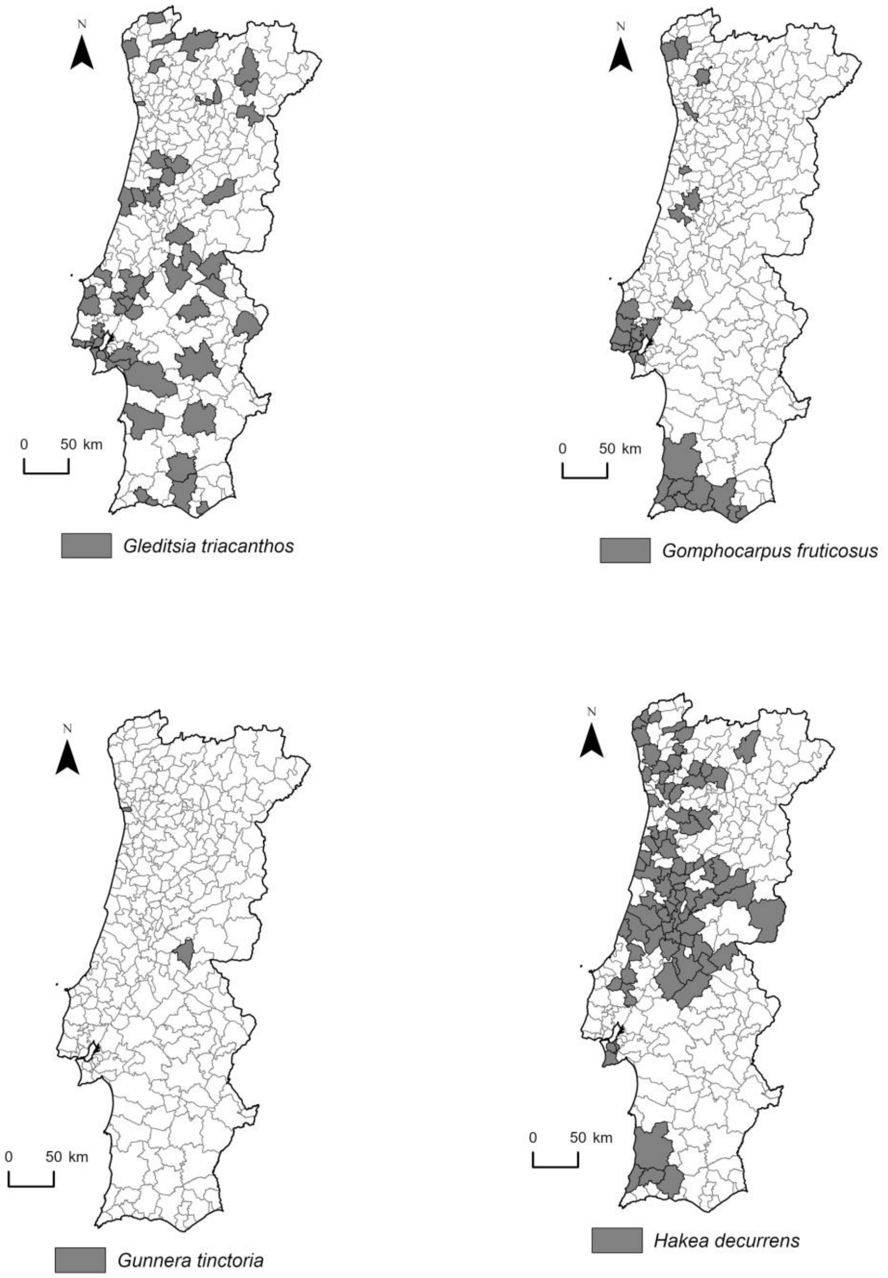
N
0 50 km
Gleditsia triacanthos
N
0 50 km
Gomphocarpus fruticosus
N
0 50 km
Gunnera tinctoria
N
0 50 km
Hakea decurrens

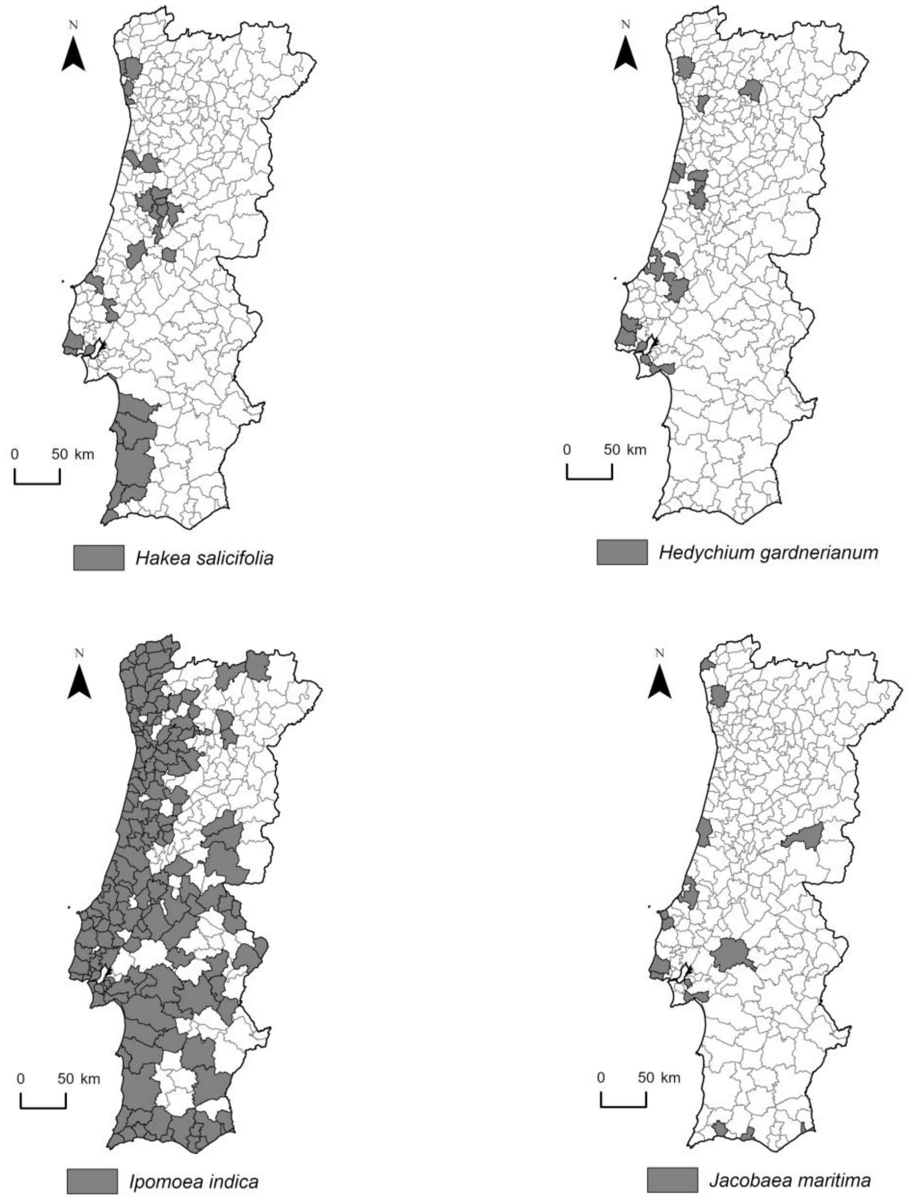
N
0 50 km
Hakea salicifolia
N
0 50 km
Hedychium gardnerianum
N
0 50 km
Ipomoea indica
N
0 50 km
Jacobaea maritima

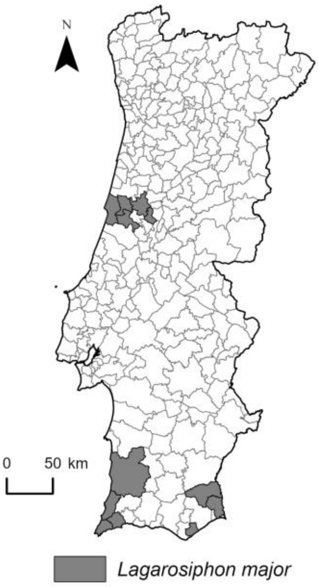
N
0 50 km
Lagarosiphon major

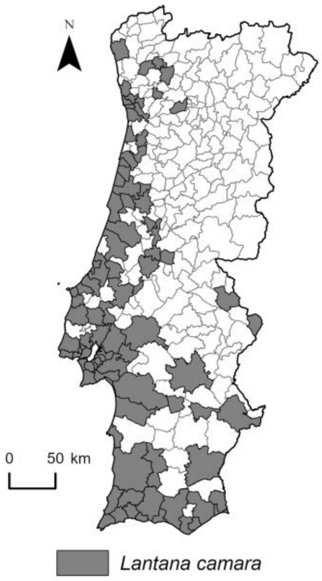
N
0 50 km
Lantana camara

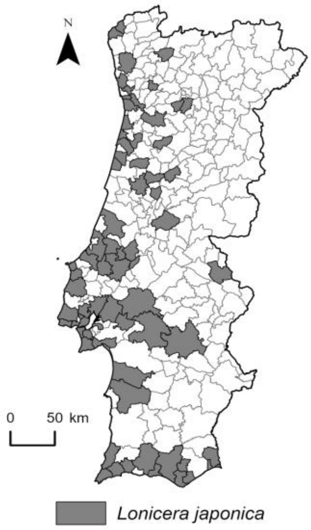
N
0 50 km
Lonicera japonica

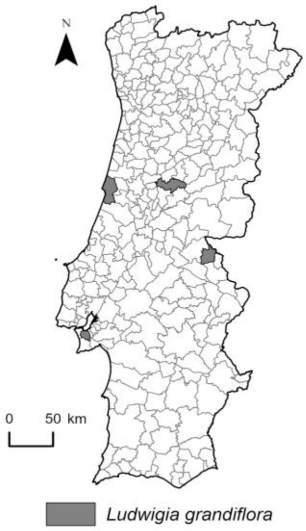
N
0 50 km
Ludwigia grandiflora

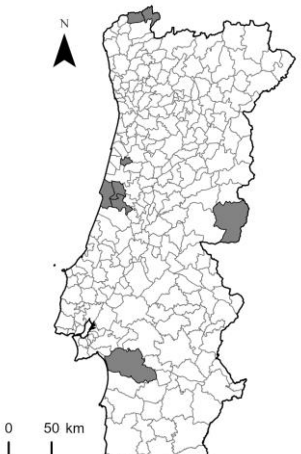
N
0 50 km
Ludwigia peploides

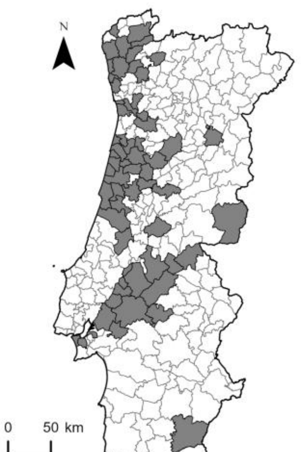
N
0 50 km
Myriophyllum aquaticum

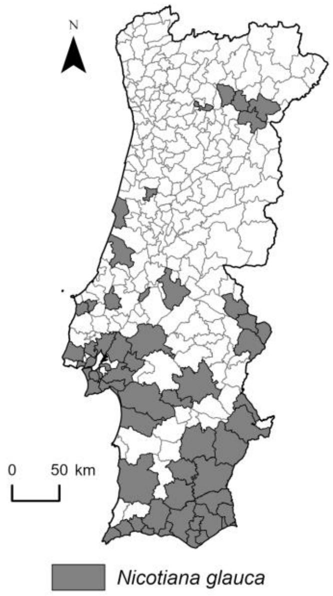
N
0 50 km
Nicotiana glauca

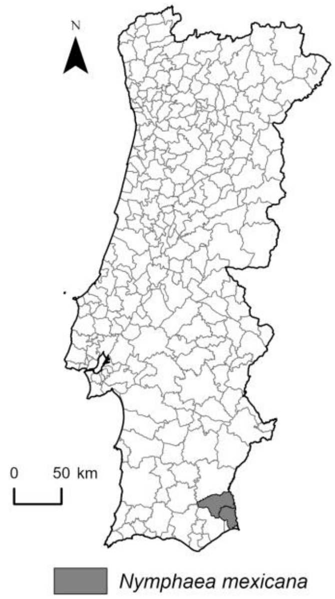
N
0 50 km
Nymphaea mexicana

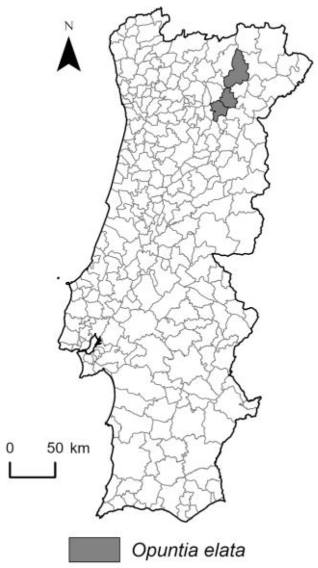
N
0 50 km
Opuntia elata

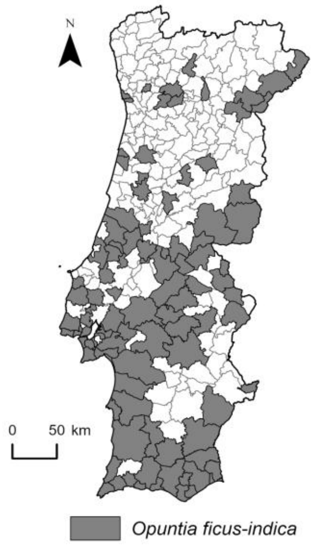
N
0 50 km
Opuntia ficus-indica

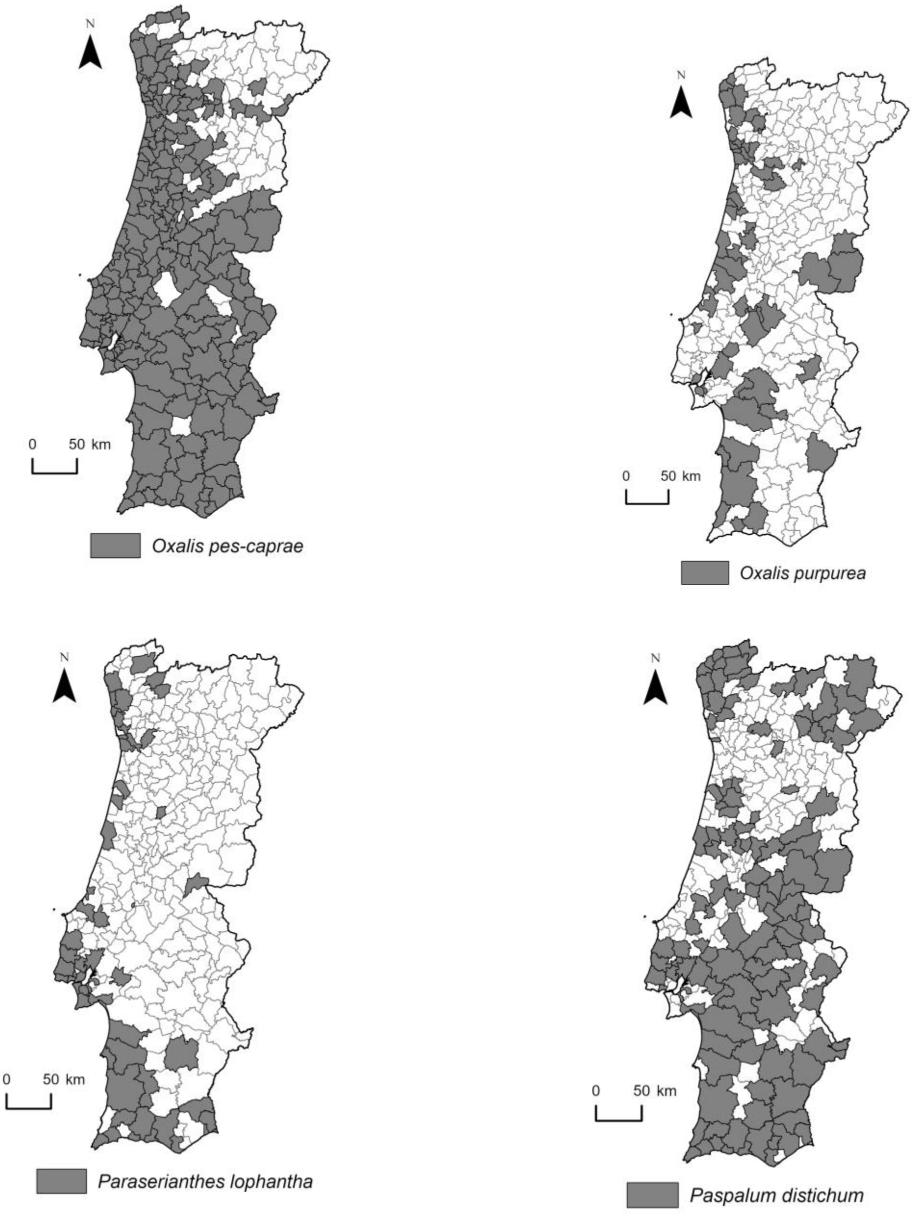
N
0 50 km
Oxalis pes-caprae
N
0 50 km
Oxalis purpurea
N
0 50 km
Paraserianthes lophantha
N
0 50 km
Paspalum distichum

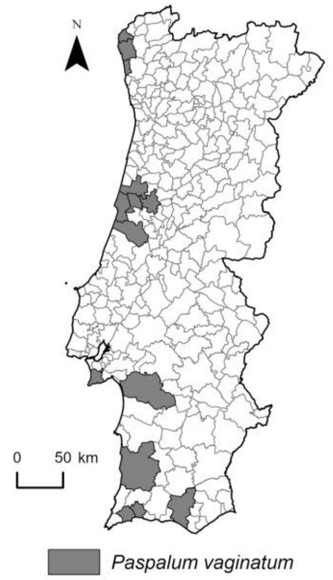
N
0 50 km
Paspalum vaginatum

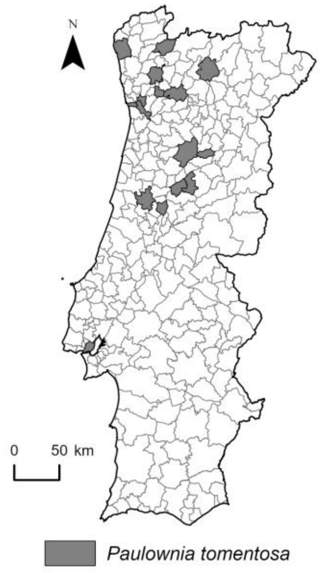
N
0 50 km
Paulownia tomentosa

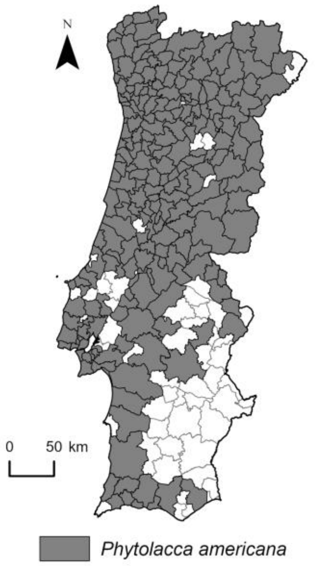
N
0 50 km
Phytolacca americana

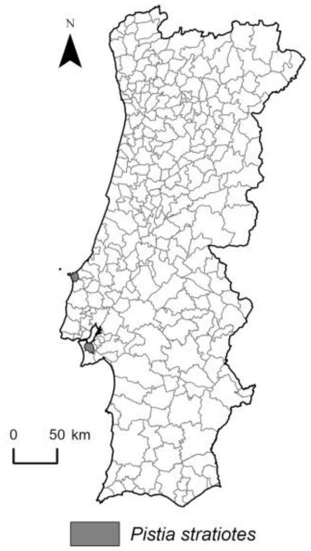
N
0 50 km
Pistia stratiotes

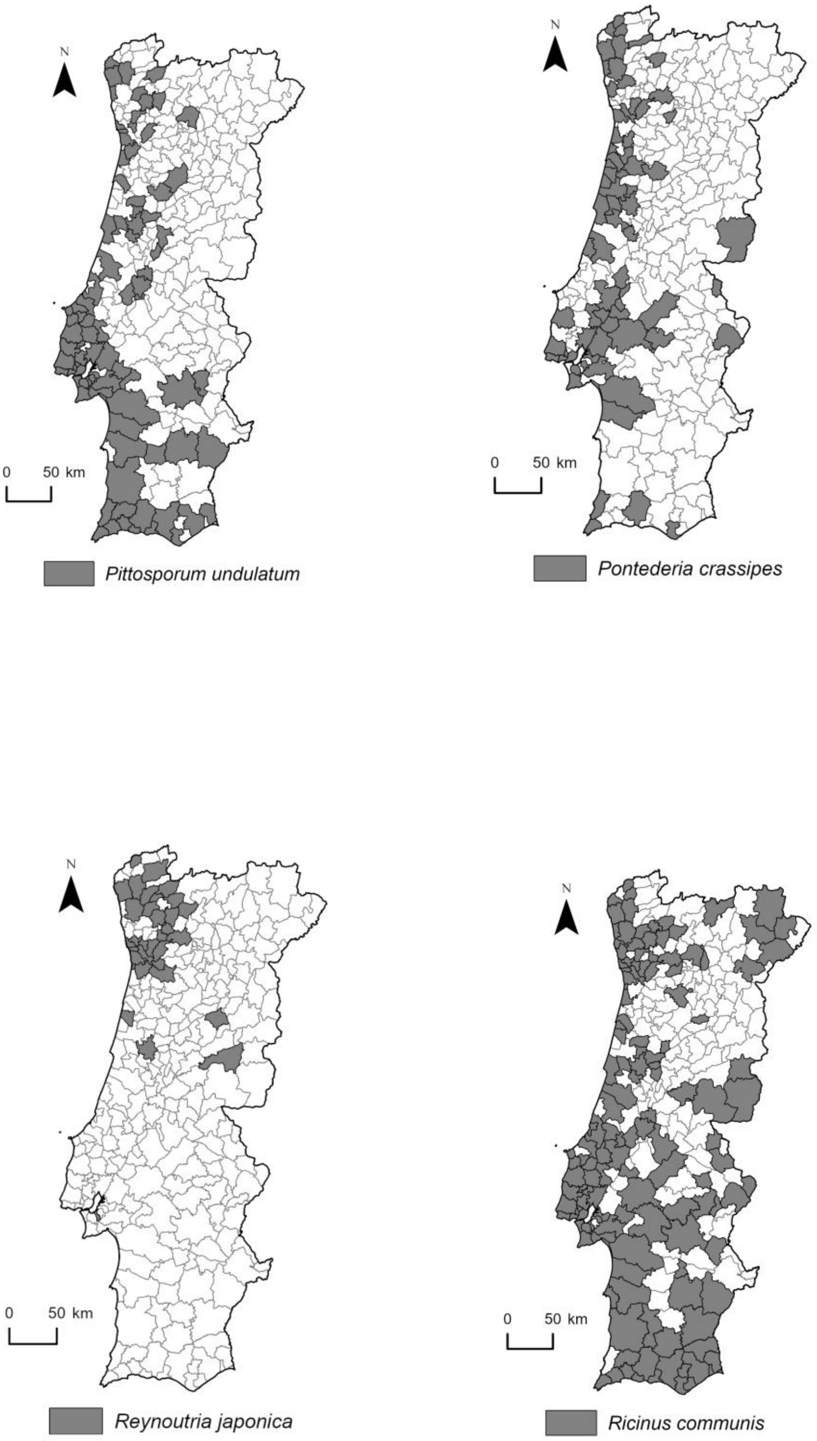
N
0 50 km
Pittosporum undulatum
N
0 50 km
Pontederia crassipes
N
0 50 km
Reynoutria japonica
N
0 50 km
Ricinus communis

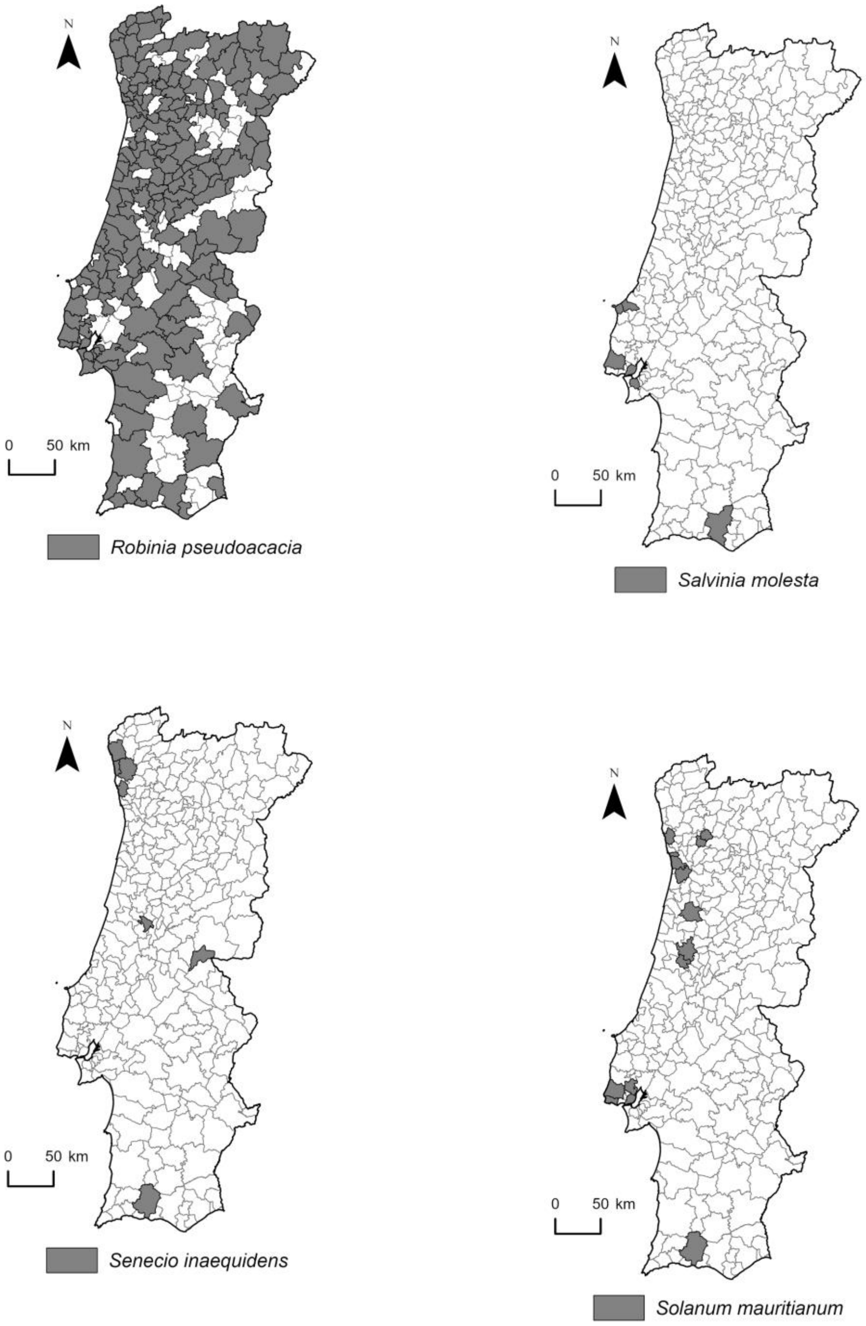
N
0 50 km
Robinia pseudoacacia
N
0 50 km
Salvinia molesta
N
0 50 km
Senecio inaequidens
N
0 50 km
Solanum mauritianum

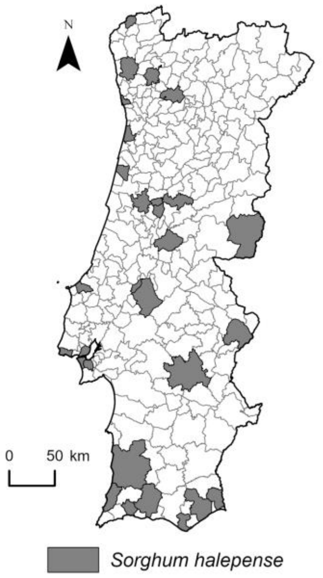
N
0 50 km
Sorghum halepense

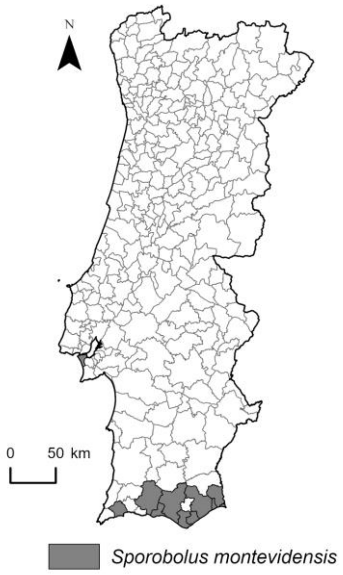
N
0 50 km
Sporobolus montevidensis

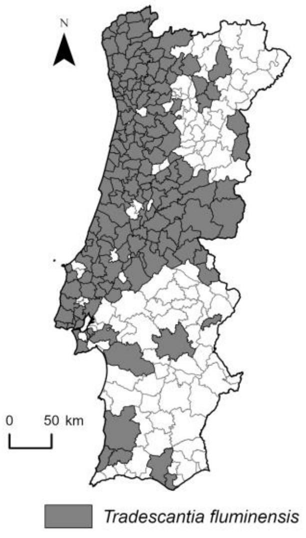
N
0 50 km
Tradescantia fluminensis

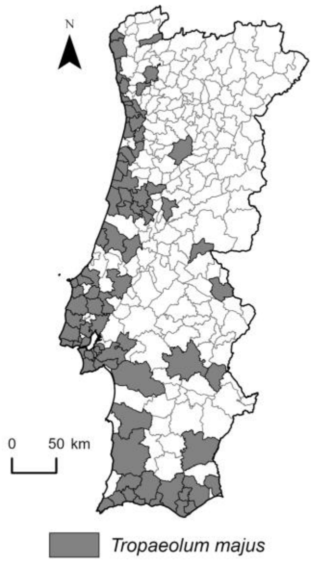
N
0 50 km
Tropaeolum majus

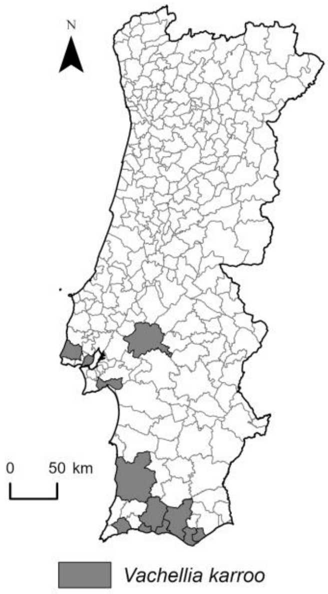
N
0
50 km
Vachellia karroo
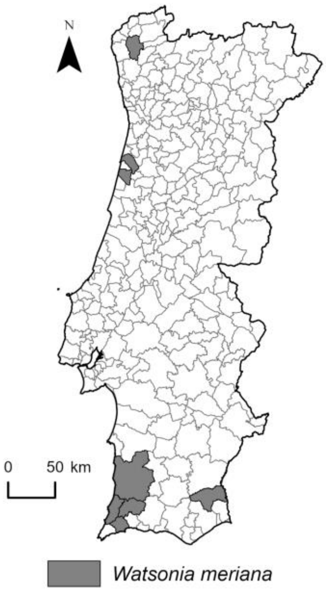
N
0
50 km
Watsonia meriana
