## Supplementary material for "Distribution patterns of invasive alien plant species in mainland Portugal": Results without FloraOn data

### Supplementary material 3

#### Results without FloraOn data

Generally, the distribution of species richness without including the Flora-On data showed similar patterns to those with the original dataset, with the municipalities located in coastal regions and major urban centers showing higher richness of invasive plant species (Suppl. material: Appendix 3, Fig. S3.1). Similarly, the groups of species followed similar distribution patterns of invasive plants along mainland Portugal (Suppl. material: Fig. S3.2). The significant positive and negative relationships with variations in species richness by municipality were identified between the same variables described above (Suppl. material: Appendix 3, Table S3.1). Nevertheless, compared with the original dataset, we observed the exclusion of one species, *Amaranthus hypochondriacus*, which had only one occurrence record from Flora-On, and the significant decrease in the number of municipalities in which the *Erigeron sumatrensis* is present, from 105 to 10.

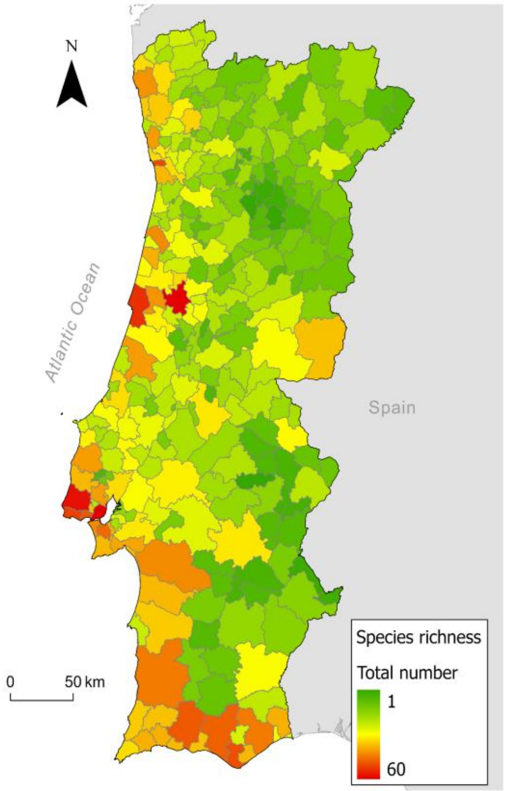


**Figure S3.1.** Total number of invasive alien plant species in the municipalities of mainland Portugal, based on the **dataset without the Flora-On occurrence records**.

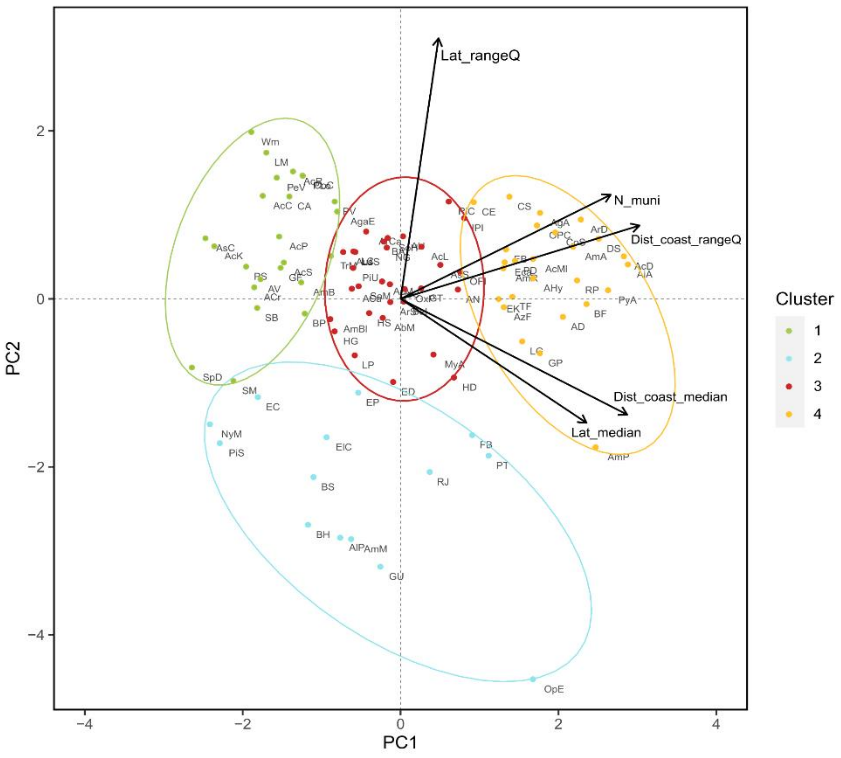


**Figure S3.2.** Groups of species resulting from the k-means clustering analysis, based on the **dataset without the Flora-On occurrence records**. Group 1 (green): species primarily located along the coast; Group 2 (blue): species with narrow ranges; Group 3 (red): moderately widespread species; Group 4 (yellow): widespread species. The PCA axes represent the total number of municipalities (N_muni), the median distances to the coast (Dist_coast_median), the interquartile range of distances to the coast (Dist_coast_rangeQ), the median latitudes (Lat_median), and the interquartile range of the latitudes (Lat_rangeQ).

**Table S3.1.** Coefficients and *p-value* results of each explanatory variable considered in the GLS model, considering the **dataset without the Flora-On occurrence records.** Significant relationships ($p < 0.05$) with variations in species richness by municipality are shown in bold.

| Explanatory variable | Coefficient (SE) | *p*-value |
|---|---|---|
| Daily mean air temperatures of the wettest quarter (°C) | -1.8857 (0.001) | 0.061 |
| Mean daily maximum air temperature of the warmest month (°C) | -1.1185 (0.031) | 0.264 |
| Ruggedness Index | 1.3096 (0.010) | 0.192 |
| Minimum distance to coastline | **-2.8968 (0.003)** | **0.004** |
| Municipality area | **8.1815 (0.006)** | **<.001** |
| Density of agricultural areas | 1.2250 (0.003) | 0.222 |
| Density of forest areas | 0.5093 (0.003) | 0.612 |
| Density of classified areas | 1.4953 (0.160) | 0.136 |
| Burnt area in the last 15 years | 0.1098 (0.002) | 0.913 |
| Soil pH | 0.7663 (1.253) | 0.541 |
| Density of local hydrographic network | 0.6702 (0.094) | 0.503 |
| Density of railways | 1.0552 (0.159) | 0.291 |
| Density of power lines | **2.6487 (0.241)** | **0.009** |
| Travel time to cities (Minutes) | **-3.2629 (0.002)** | **0.001** |
| Sampling effort | 1.7271 (0.025) | 0.086 |
| Number of iNaturalist projects with IAP records | 1.7217 (0.025) | 0.086 |
| Number plant nurseries and aquarophilia stores | 1.3346 (0.046) | 0.183 |
